# Export dynamics of protists across the southern subtropical frontal zone reveal taxon-specific patterns

**DOI:** 10.1101/2025.02.06.636886

**Authors:** Denise Rui Ying Ong, Andrés Gutiérrez-Rodríguez, Jaret Bilewitch, Scott Nodder, Michael R. Stukel, Moira Décima, Adriana Lopes dos Santos

## Abstract

Gravitational particle sinking is the main mechanism for carbon export in the biological carbon pump. However, the export dynamics of the particle-associated protist community are not fully understood. We used 18S rRNA gene metabarcoding to characterise the exported protist community within sinking particles and bathypelagic surficial sediments in oligotrophic subtropical and high-nutrient, low-chlorophyll subantarctic waters. Sinking particles were collected with formalin-fixed and preservative-free particle interceptor traps (fixed and live traps, respectively) to identify the community involved in particle export (fixed) and protist loss from remineralisation (live). We paired this with community analysis of the upper and lower water column (mixed layer and below mixed layer to mesopelagic, respectively) to compare the relative sources of exported protists. Amplicon sequences variants (ASVs) from upper water column samples accounted for 2 to 4-fold higher proportion of reads and ASV rich-ness compared to lower water column samples in fixed trap and sediment samples, suggesting low influence of the suspended protist community from the lower water column on export. We further traced the export patterns of upper water column protist taxa by analysing the change in taxa relative abundance across the mixed layer to mesopelagic depths. Export patterns differed between taxa, which is similarly suggested by taxa-specific loss of ASV richness between fixed and live traps, but remained the same across biogeochemically-contrasting water masses. This could imply that the drivers for protist loss during export are related to characteristics consistent across environmental conditions, such as specific microbial interactions or inherent cell properties.

## Introduction

The biological carbon pump is an important process by which a fraction of photosynthetically fixed carbon at the surface ocean is moved to the deep ocean, and sequestered up to geological time scales (De La Rocha and Passow 2007). The main way particulate organic carbon (POC) is moved to the deep ocean is through the sinking of particles made up of cells that are repackaged into aggregates, detritus or faecal pellets, as well as individual living and dead cells (Durkin et al. 2021; Nowicki et al. 2022; Ricour et al. 2023).

Despite the crucial role of the biological carbon pump for the functioning of marine ecosystems and as a buffer for climate change (Gruber et al. 2019; Sabine et al. 2004), yearly estimates of global carbon export are poorly constrained (Henson et al. 2022; Ricour et al. 2023), and vary across space (Martin et al. 1987; Weber et al. 2016) and time (Giering et al. 2017). Part of this variation is due to the complex influence of microbial community composition and activity in sinking particles (Burd et al. 2010). As the particles sink, they are subjected to microbial community processes of consumption, fragmentation, and remineralisation within and outside the particles (Cavan et al. 2017; De La Rocha and Passow 2007; Giering et al. 2014). This transformation further influences the remineralisation depth of the sinking particles (Kwon et al. 2009) and affects the efficiency of carbon export, raising the need to understand community interactions within the sinking particles (Buesseler et al. 2007; Burd et al. 2010; Worden et al. 2015). Importantly, the surface community influences the community composition present in the sinking particles and the types of particles formed, and as a consequence sinking speeds and remineralisation rates (Bach et al. 2019; Guidi et al. 2009; Weber et al. 2016), underlining that the two parts of the biological carbon pump (particle export from the surface ocean across the mixed layer, and transfer to mesopelagic and bathypelagic depths) to are not mutually exclusive and have to be considered together.

Sinking particles are sampled through the deployment of particle interceptor traps (PITs; Knauer et al. 1979), and its community composition can be characterised using microscopy and pigment analysis (Caron et al. 1986; Mackinson et al. 2015; Wilks et al. 2021), or more recently using DNA sequencing to target the prokaryotic (LeCleir et al. 2014; Li et al. 2023) and eukaryotic (Amacher et al. 2009; Durkin et al. 2022) communities. However, assessing the contribution of microbial communities toward export to deep ocean sediments is not complete with this approach, as microbial communities present in PIT- sampled particles might not be deposited in the sediments. A varying proportion of genetic material from pelagic protist (or microbial eukaryotic) taxa have been detected in the sediments (Cordier et al. 2022; Laroche et al. 2020; Morard et al. 2017; Nguyen et al. 2024; Pawlowski et al. 2011), presumably transported via sinking particles from the surface of the ocean. Yet, the connectivity between surface and sinking particles (Amacher et al. 2009; Durkin et al. 2022; Gutierrez-Rodriguez et al. 2019), as well as between sinking particles and sediments (Preston et al. 2020), have only been explored in separate studies. Additionally, suspended prokaryotes (Li et al. 2023) and protists (Fontanez et al. 2015; Gutierrez-Rodriguez et al. 2019) residing below the mixed layer can colonise the sinking particles and transform the particle-associated microbial community or form new particles (Bochdansky et al. 2017; Huffard et al. 2020; Nguyen et al. 2022). While suspended prokaryotic communities from the lower water column can have significant contribution toward particle-associated communities compared to the upper water column (Li et al. 2023), the relative contribution of the upper and lower water column protists within sinking particles is unknown.

The study was conducted in biogeochemically-distinct subtropical and subantarctic water masses across the southern subtropical frontal zone, east of Aotearoa-New Zealand in the southwest Pacific Ocean during the austral spring. Northern subtropical waters are warm and mainly nitrogen-limited (Zentara and Kamykowski 1981), while southern subantarctic waters are cool with high-nutrient, low-chlorophyll (HNLC) characteristics (Boyd et al. 1999). The contrasting surface protist communities in subtropical and subantarctic waters (Gutiérrez-Rodríguez et al. 2022) make it an ideal sampling location to explore the effects of surface community on export.

We aimed to address these research gaps by characterising the particle-associated protist community from the surface mixed layer to bathypelagic sediments during the same sampling cruise, to determine the sources of the exported community, and how they evolve from the surface to deep ocean sediments. We applied DNA metabarcoding of the 18S rRNA gene to characterise the taxonomic composition of formalin-fixed (fixed traps) and preservative-free (live traps) PITs deployed from below the mixed layer to mesopelagic depths, where microbial activity and DNA degradation were inhibited in fixed traps but allowed to continue in live traps (Fontanez et al. 2015; Gutierrez-Rodriguez et al. 2019). We also sampled the upper (mixed layer; 5–40 m) and lower (below mixed layer to mesopelagic; 70–500 m) water column, as well as the bathypelagic surficial sediments (428–2604 m). Our specific objectives were to assess the relative contribution of the suspended protist communities from the upper and lower water column on exported communities in sinking particles and sediments, and the changes in relative abundance of the particle-associated protist taxa during export from the surface to mesopelagic depths. We further examined taxa loss in particles from remineralisation by comparing the communities in fixed and live traps. To our knowledge, this is the first study directly connecting the particle-exported protist community at amplicon sequence variant (ASV) level (i.e., DNA sequence level) from the surface mixed layer to bathypelagic sediments.

## Material and Methods

### Study area and sampling

Sampling was carried out during the Salp Particle expOrt and Oceanic Production (SalpPOOP) cruise on board the R/V Tangaroa (TAN1810, 21st October – 21st November 2018), near the Chatham Rise, east of Aotearoa-New Zealand. The sampling strategy was designed based on Lagrangian experiments called “cycles”, where the same water parcel was followed over multiple days using a satellite-tracked floating array with a drogue centred at 15 m depth (Landry et al. 2009). Water column profiles of temperature, salinity, fluorescence and photosynthetically active radiation (PAR) were obtained with Seabird (SBE 911plus) CTD (Conductivity-Temperature-Depth) attached to a rosette frame with 10 L Niskin bottles for surface water collection. Each cycle had a duration of 3 to 7.5 days, with two in subtropical waters (ST1 and ST2) and three in subantarctic waters (SA-Sc, SA1 and SA2; Figure 1). SA-Sc lasted for 7.5 days and the cycle was divided into two periods of 5 (A) and 2.5 days (B), each with distinct nutrient, phytoplankton biomass and productivity measurements (Décima et al. 2023). Measurements and samples were collected from both periods of SA-Sc, unless otherwise stated.

**Figure 1:**
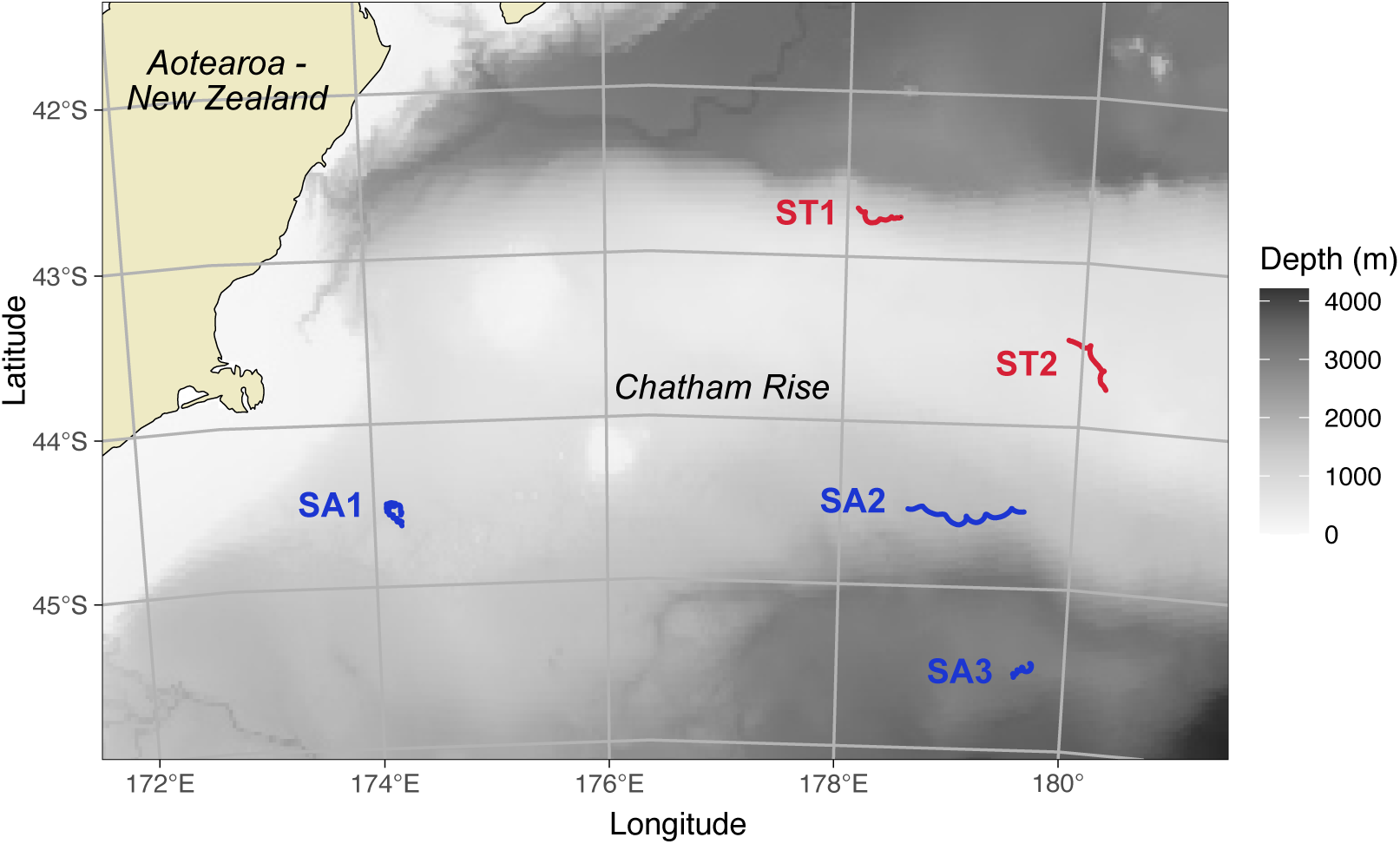
Map of the study area. The path followed during each sediment trap deployment is indicated in red for subtropical (ST) cycles and blue for subantarctic (SA) cycles.

Surface seawater samples were obtained from daily pre-dawn CTDs (0200h deployment) to conduct standard ^14^C-fixation experiments for net primary productivity (NPP) measurements, and from morning CTDs (0900-1200h deployment) to measure nitrate and size-fractionated Chlorophyll *a* (Chl *a*) concentration. These samples were collected from several depths (5 to 70 m; Table S1) and processed as described in Décima et al. (2023).

Seawater and sediment samples were collected for DNA sequencing. Seawater samples from the upper water column were collected using Niskin bottles (0200h deployment) twice each cycle from three to six depths (5 to 40 m; Table S1), corresponding to the mixed layer as estimated by CTD profiles. 1.4–2.4 L of seawater from each Niskin bottle was filtered through a 0.22 µm pore-size Sterivex filter (Millipore) using a peristaltic pump and flash frozen in liquid nitrogen. Seawater samples from the lower water column were collected using high-volume in situ McLane pumps from four depths (70 to 500 m), starting from 30 m below the mixed layer depth to mesopelagic depths. The McLane pumps were deployed once per cycle (only in SA-Sc-B of cycle SA-Sc) at the same depths as the particle interceptor traps (PITs), except for SA2 (Table 1). The pumps filtered for 120–150 minutes and collected 200 L of water onto polycarbonate 0.2µm filters. Surficial sediment samples were collected with an Ocean Instruments MC-800 multicorer once each cycle (only in SA-Sc-a of cycle SA-Sc; Table 1), where a surface scrape removed 25 mL of sediment into a 50 mL Falcon tube (see Supplementary Material). All samples were stored at −80 ^◦^C until processing.

**Table 1:** Depths (m) sampled of each sample type across the cycles for DNA metabarcoding. The upper water column was sampled with a CTD across 3–6 depths between the ranges specified. Fixed and live sediment traps were deployed at 4 depths below the deep chlorophyll maximum at the depths reported. At the same depths, the lower water column was sampled with a large volume McLane pump, except for SA2 where the depths sampled of McLane pump (in brackets) is 40 m higher than what is reported for sediment traps.

| Sample type | Depth level | ST1 | ST2 | SA-Sc | SA1 | SA2 |
| --- | --- | --- | --- | --- | --- | --- |
|  |  | Depth (m) |  |  |  |  |
| Upper water column (Niskin) |  | 5–40 | 5–40 | 5–40 | 5–40 | 5–30 |
| Fixed traps, live traps,<br>lower water column<br>(McLane pump) | 1 | 70 | 70 | 70 | 70 | 110 (70) |
|  | 2 | 100 | 100 | 100 | 100 | 140 (100) |
|  | 3 | 300 | 200 | 300 | 300 | 340 (300) |
|  | 4 | 500 | 300 | 500 | 500 | 540 (500) |
| Sediment |  | 1051 | 428 | 780 | 1405 | 2604 |

### Particle Interceptor Traps

Particle interceptor traps (PITs) were used to collect sinking particles for DNA sequencing and assess export fluxes of particulate organic carbon (POC), Chl *a* and phaeopigment concentration. The VERTEX style PITs (Knauer et al. 1979) were set up with free-floating surface-tethered arrays, equipped with a surface Iridium beacon and light flasher, surface and sub-surface floats, 10 m-long bungies to reduce wave action and holey-sock drogue at 15 m. They were deployed once at the start of each experimental cycle (only in SA-Sc-a of cycle SA-Sc) for 2.8 to 4 days at four depths (Table 2), typically starting from 30 m below the mixed layer depth (estimated by CTD profiles, 70 m for almost all the cycles), as well as 100 m, 300 m and 500 m below the surface (Table 1). The array consisted of 12 PIT cylindrical traps (inner diameter: 7 cm and length: 58 cm) on each cross-frame, with one cross-frame for each depth. Prior to deployment, each cylinder was filled with a high density brine solution to just below the bottom of the baffles, with or without formalin (0.4% formaldehyde final concentration) for fixed (DNA, POC, Chl *a* and phaeopigment) and live (DNA) traps, respectively. There were no live trap samples collected for SA-Sc. Samples for DNA sequencing were filtered to obtain the total (*>* 0.2µm) and size fractionated (pico-nano: 0.2-20 µm and micro: *>* 20µm) community for both fixed and live traps (Figure S1). The community within fixed and live trap samples were averaged across all size fractions for all analysis. Details are provided in the Supplementary Material.

**Table 2:** Physical, chemical and biological properties of each cycle, as well as sediment trap deployment dates and duration. Cycle SA-Sc was further divided into two time periods of “SA-Sc-a” and “SA-Sc-b”, where sediment traps deployed to collect sinking particles for DNA sequencing was only conducted during “SA-Sc-a”, while McLane pump was deployed in “SA-Sc-b”. Cycles are listed in chronological order.

| Cycle | Surface salinity | Surface temp (°C) | NO<sub>3</sub> (µm) | NPP (mgC m<sup>-2</sup> day<sup>-1</sup>) | Chl *a* (mg m<sup>-2</sup>) | Dates deployed | Duration (Days) |
| --- | --- | --- | --- | --- | --- | --- | --- |
| SA-Sc-a | 34.58 ± 0.02 | 10.74 ± 0.18 | 1.68 ± 0.32 | 638.2 ± 235.6 | 73.1 ± 20.3 | 24–28 Oct 2018 | 3.43 |
| SA-Sc-b | 34.61 ± 0 | 11.35 ± 0.05 | 1.26 ± 0.13 | 389.9 ± 5.3 | 51.8 ± 9 |  |  |
| SA1 | 34.37 ± 0.01 | 10.20 ± 0.07 | 2.27 ± 0.07 | 321.4 ± 62.2 | 37.5 ± 5.5 | 2–6 Nov 2018 | 4.07 |
| ST1 | 34.98 ± 0.01 | 13.38 ± 0.07 | 0.33 ± 0.1 | 755.5 ± 371.5 | 74.2 ± 20.1 | 7–10 Nov 2018 | 2.89 |
| ST2 | 34.75 ± 0.02 | 12.99 ± 0.41 | 0.62 ± 0.12 | 452.3 ± 102.1 | 37.6 ± 3.3 | 12–15 Nov 2018 | 2.96 |
| SA2 | 34.36 ± 0 | 11.40 ± 0.02 | 2.24 ± 0.02 | 241.2 ± 31.9 | 20.2 ± 2.7 | 16–19 Nov 2018 | 2.93 |

### DNA extraction, PCR and amplicon analysis

DNA was extracted from the upper water column samples using the DNeasy mini Blood and Tissue Kit (Qiagen, Germany), lower water column and PIT samples using the Nucleospin Plant II DNA extraction kit (Macherey-Nagel), and sediment samples using the DNeasy mini Powersoil Kit (Qiagen, Germany), with modified protocols. The V4 region of the 18S rRNA gene of eukaryotes was amplified with the protocol described in Décima et al. (2023), using the primer set TAReuk454FWD1 (5’- CCAGCASCYGCGGTAATTCC-3’) and V4 18S Next. Rev (5’-ACTTTCGTTCTTGATYRATGA-3’; Piredda et al. 2017). Samples were purified, barcoded, and sequenced using the Illumina MiSeq platform (2 *×* 250 bp; INRA Auzeville, France). Sequence data was processed on RStudio Version 1.4.1717 (RStudio Team 2021) using DADA2 R package Version 1.12 (Callahan et al. 2016) and taxonomy was assigned against PR2 database version 4.14 (Guillou et al. 2013). Details are provided in the Supplementary Material.

## Results

### Upper water column characteristics and export flux

Subtropical cycles exhibited warmer and more saline conditions compared to subantarctic cycles, with lower nitrate concentrations (Table 2). SA-Sc had intermediate temperature, salinity and nitrate concentrations, likely reflecting the influence of the Southland Current, which is composed of mainly subantarctic waters mixed with neritic subtropical waters. Surface Chl *a* concentration and NPP were on average higher for subtropical compared to subantarctic cycles (Table 2).

POC fluxes were highest in cycles closest to the subtropical front (ST2, SA1 and SA-Sc; Figure 2A). The overall low Chl *a* to phaeopigment ratio suggest that the traps were dominated by degraded material (Figure 2B).

**Figure 2:**
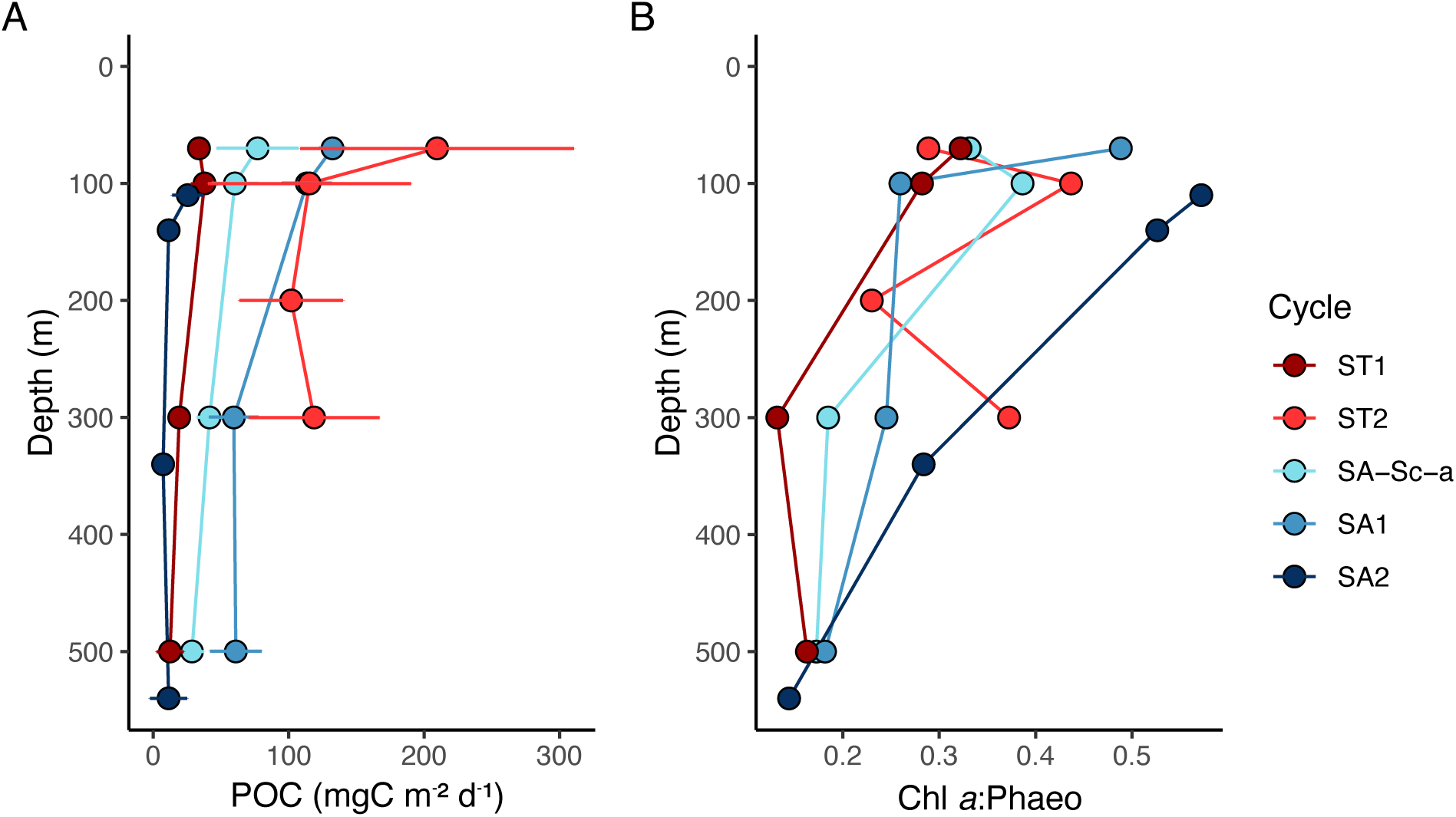
Fluxes of (A) particulate organic carbon (POC) and (B) Chlorophyll *a* to phaeopigment ratio in fixed sediment traps. Colours represent the cycles. Error bars represent the standard deviation of POC fluxes (n = 3).

### Diversity and community distribution patterns between sample types

Across the five different sample types (upper water column, lower water column, fixed traps, live traps and sediments), we obtained 5,104 eukaryotic ASVs, of which 4,449 ASVs were assigned to protist taxa, 653 ASVs to Metazoans and 2 ASVs to subdivision Ichthyosporea. Protisan reads (68.3%) dominated the overall community over Metazoans (31.7%). Samples from the lower water column (sampled with McLane pump), fixed traps and upper water column (sampled with Niskin bottles) and had the highest number of unique protist ASVs, followed by samples from live traps and sediment samples (Figure 3B). Despite sediment samples having the lowest total number of ASVs, the average alpha diversity and ASV richness was highest in samples from sediment and upper water column, followed by lower water column and fixed traps, and finally by live trap samples (Figure S2). Alpha diversity and ASV richness measurements were similar between cycles for each sample type (Figure S3).

**Figure 3:**
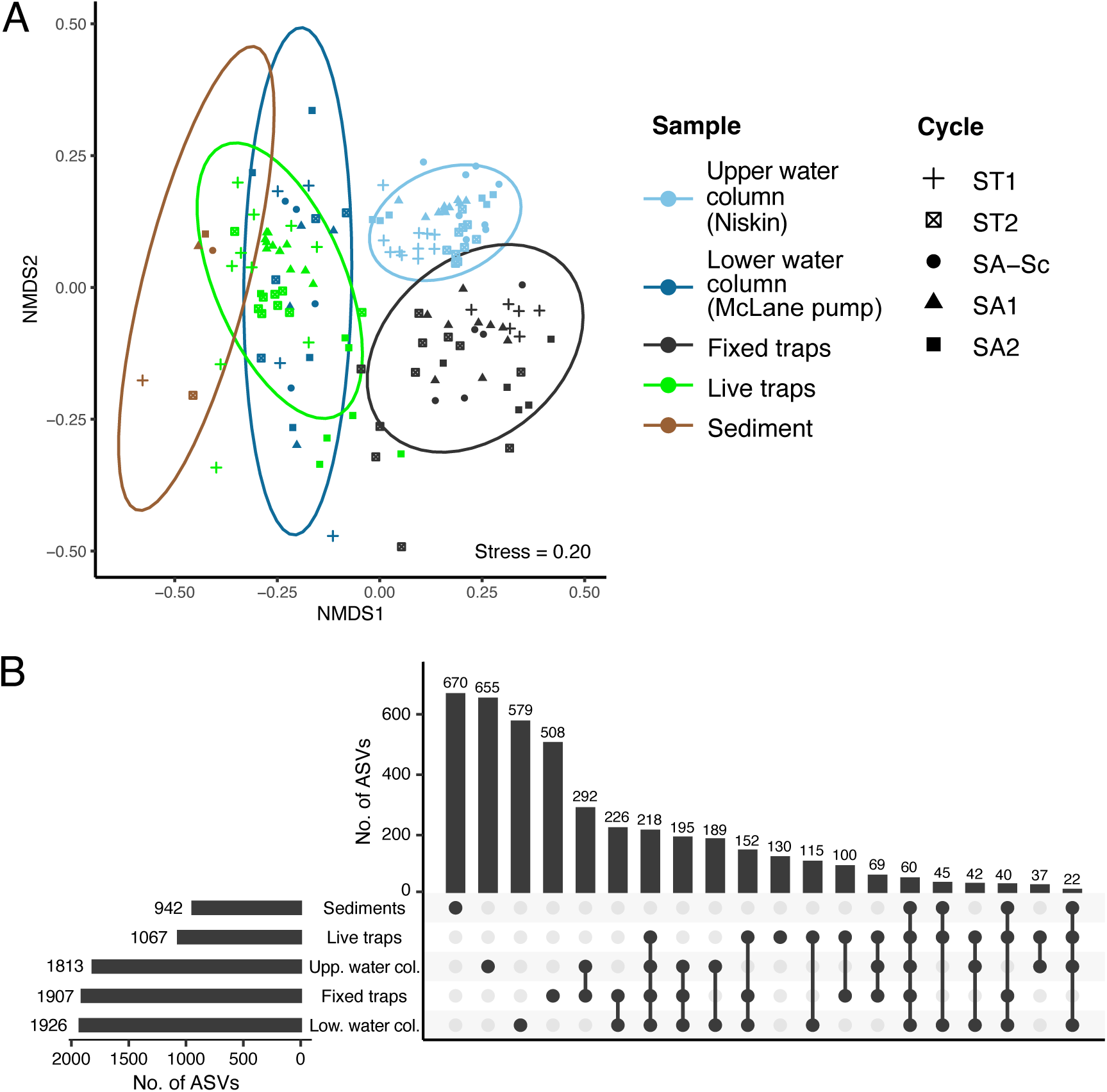
(A) Non-parametric multidimensional scaling ordination (nMDS) plot based on Bray–Curtis dissimilarity based on protist ASV composition. Sample type is indicated by colour, and cycle is indicated by shape. 95% confidence ellipses for each sample type are represented. (B) Protist ASVs common between one or more sample types (UpSet plot).

Community composition was significantly different between all sample types (Figure 3A; PERMANOVA-Adonis: R^2^ = 0.24, *P* < 0.001; Pairwise analysis: adjusted *P* < 0.01 between all groups) and water mass (PERMANOVA-Adonis; R^2^ = 0.04, *P* < 0.001), although water mass had a lesser influence than sample type. Fifty-seven percent of total ASVs (2,542 out of 4,449 ASVs) were restricted to a single sample type (Figure 3B). Sediment samples were the most distinct with 71% of ASVs that were unique to the sample type, while live trap samples were the least distinct with 88% of its ASVs shared with another sample type (Figure 3B).

On average, Dinoflagellata was the most abundant protist group (52.3*±*18.3%), followed by Radiolaria (13.7*±*20.3%), Ciliophora (8.0*±*11.8%), Chlorophyta (7.3*±*11.1%) and Gyrista (6.4*±*6.3%). The proportion of these dominant groups varied between sample types and cycles (Figures 4 and 5). A detailed description of the eukaryotic composition in upper water column, lower water column and sediment samples is included in the Supplementary Information.

**Figure 4:**
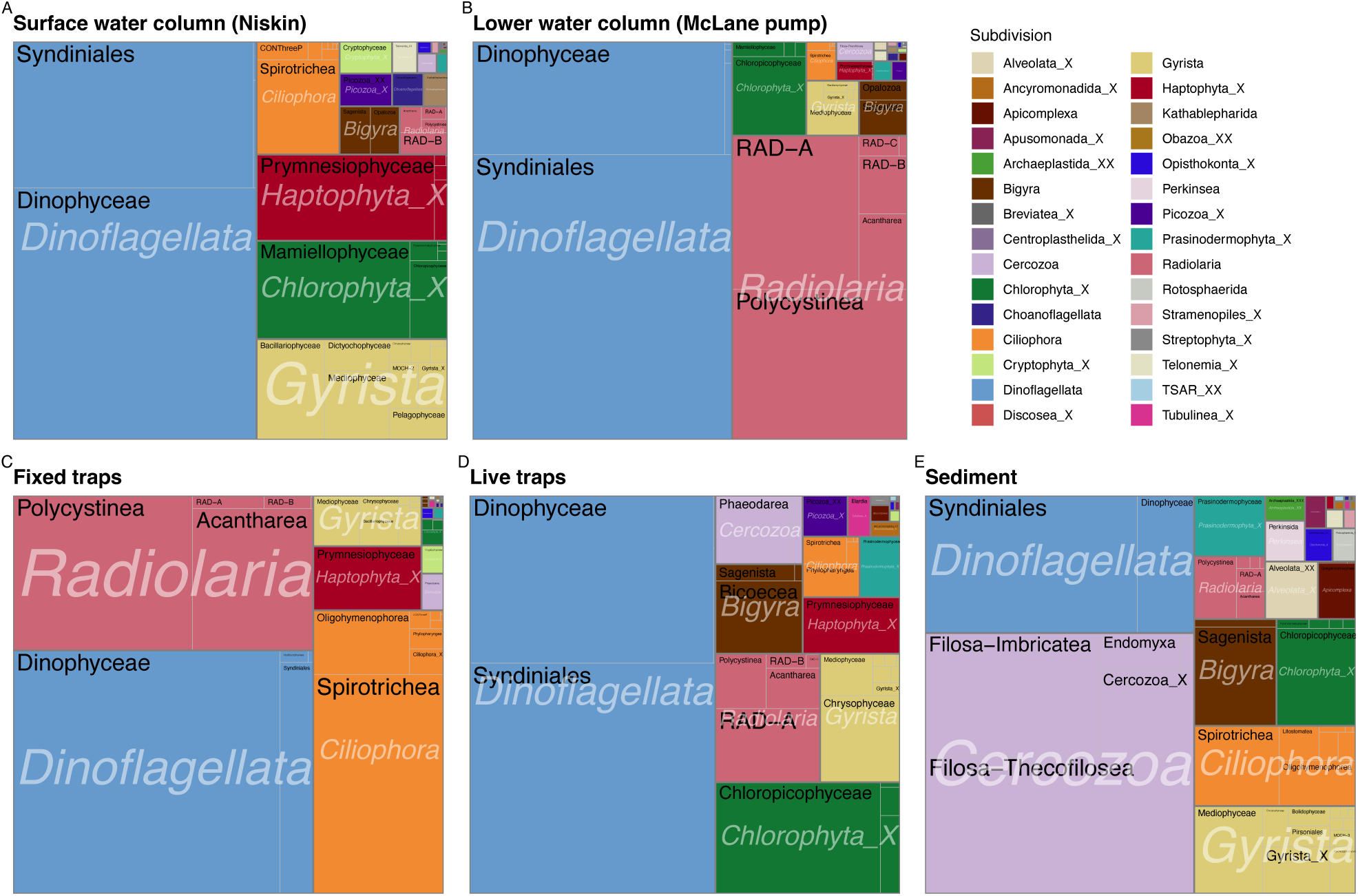
Treemap of 18S V4 rRNA protist community at division (white italics) and class (black) level, grouped by sample type. Rectangle surfaces within each treemap are proportional to the relative abundance of each taxa.

**Figure 5:**
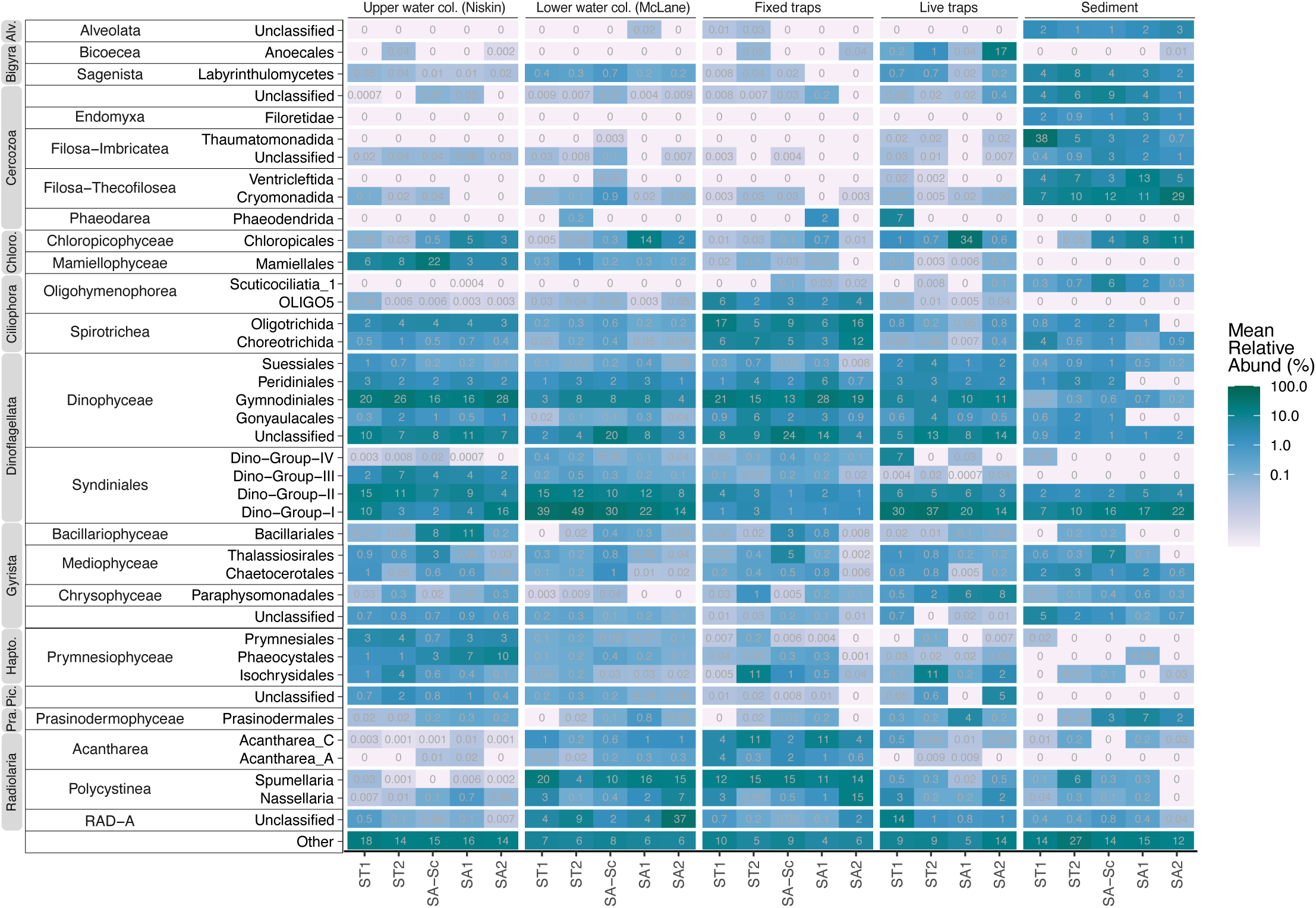
Heatmap showing the relative mean abundance (%) of 18S V4 rRNA protist reads in each cycle at order level, grouped by sample type and cycle. Samples are ordered from left to right across a spatial gradient, from subtropical (ST) to subantarctic (SA) cycles. Taxa are grouped by division and class. Division levels which are abbreviated are Alveolates (Alv.), Chlorophyta (Chloro.), Haptophyta (Hapto.), Picozoa (Pic.) and Prasinodermophyta (Pra.). ‘Other’ represents all other taxa with a mean relative abundance of less than 3% in all cycles.

### Protist composition in fixed traps

In the fixed trap samples, Dinoflagellata (42.6*±*17.1%), Radiolaria (27.1*±*24.6%) and Ciliophora (21.5*±*16.9%) were the main subdivisions present across all cycles (Figure 4C). Dinophyceae had an 8-fold higher relative abundance than Syndiniales (37.9*±*15.7% and 4.5*±*3.1%, respectively; Figure 5). Radiolaria were mostly composed of Spumellaria (13.3*±*23.6%) and Acantharia clade C (7.5*±*12.3%). Ciliophora comprised of Spirotrichea (Choreotrichida and Oligotrichida) and Oligohymenophorea.

A minor proportion of reads were assigned to photosynthetic taxa, with Isochrysidales and Thalassiosirales having higher relative abundance. *Gephyrocapsa huxleyi* (Isochrysidales) represented up to 28% of reads in ST2 (Figure S4), while *Minidiscus variabilis* (Thalassiosirales) had a higher proportion of reads in SA-Sc (5.9%; Figure 5). Other photosynthetic taxa present at very low abundance (< 1%) include Mamiellophyceae, Chloropicophyceae (mostly *Chloroparvula pacifica*), Bacillariophyceae, Chrysophyceae and Prasinodermophyceae (only *Prasinoderma singularis*).

### Sources and loss of particle-exported protist community

We investigated the upper (mixed layer) and lower (below mixed layer to mesopelagic depths) water column as sources of the exported protist community in sinking particles and sediments. Here, we identified protist ASVs in the fixed trap and sediment samples that were common with the upper or lower water column (referred to as “shared ASVs”) separately for each water mass (subtropical and subantarctic). ASVs present in both upper and lower water column samples were classified as from the upper water column, assuming that they were present in the lower water column via vertical export. To justify this assumption, we analysed the community composition of ASVs present in all three sample types (fixed traps, upper and lower water column). Most of these shared ASVs had a higher relative abundance in upper compared to lower water column samples, except Radiolaria and Syndiniales (Figure S5), which were also dominant in the lower water column samples (Figure 4B). As the community was dominated by Dinoflagellata across all sample types (Figure 4), Dinoflagellata ASVs were analysed and presented separately below.

Overall, shared ASVs from the upper and lower water column represented 81.2% of reads and 57.7% of ASVs in fixed trap samples, and 22.1% of reads and 13.8% of ASVs in sediment samples. Shared ASVs from the upper water column accounted for a 2 to 4-fold higher relative abundance compared to lower water column in fixed traps (average from upper: 52.4%, lower: 28.9%) and sediment samples (upper: 17.5%, lower: 4.6%; Figure 6A), and nearly twice the number of ASVs (fixed traps: 355 and 204, sediment: 40 and 26 ASVs from the upper and lower water column, respectively; Table S2). Across the depths sampled, the relative contribution of shared ASVs in fixed traps varied slightly, with no obvious trend of increase or decrease (Figure S6). Protist ASVs shared between the upper water column and fixed traps encompassed 45% of ASVs and 72.7% of reads in the upper water column, indicating that most of the community, i.e. likely the more abundant taxa, could be exported from the surface through particle sinking. Yet, 90% of shared ASVs corresponding to three-quarters of reads in fixed traps were not detected in the sediments, suggesting that a large proportion of exported protist diversity was lost during particle sinking to the sediments. There was also a difference in the sources of shared ASVs found in sediment samples between the water masses. In subtropical sediments, upper water column ASVs represented similar relative abundance compared to lower water column ASVs (8.3*±*5.1% and 7.2*±*1.2%, respectively; Figure 6A). In subantarctic sediments, upper water column ASVs represented 8-fold higher relative abundance compared to lower water column ASVs (23.6*±*3.4% and 2.8*±*2.2%, respectively).

**Figure 6:**
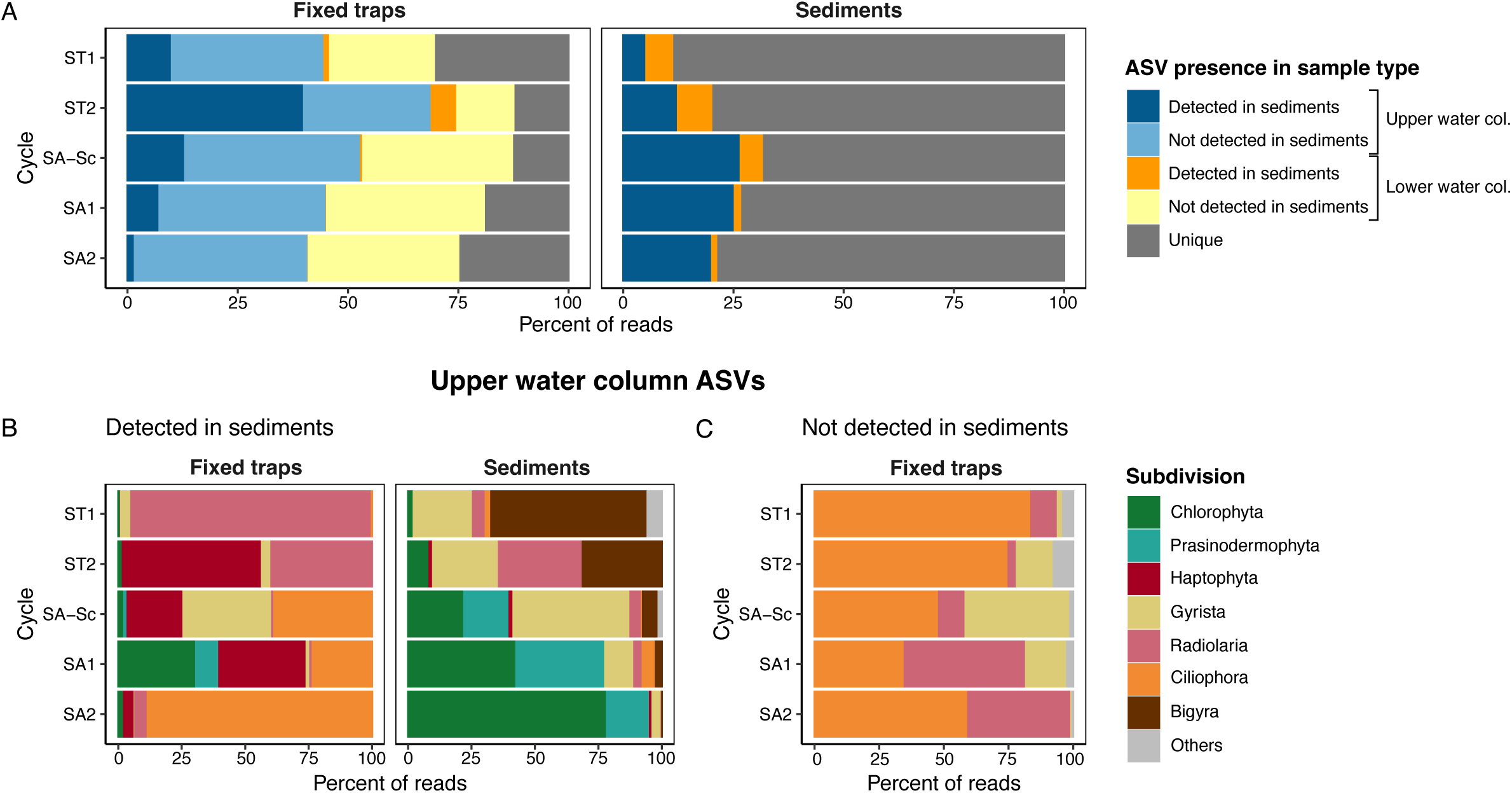
Protist community (excluding Dinoflagellata) from the fixed traps and sediment that was common between the upper (mixed layer, sampled with Niskin) and lower (below mixed layer to mesopelagic depths, sampled with McLane pump) water column. A) Protist ASVs in fixed traps and sediment samples were characterised as from the upper or lower water column, further divided into ASVs that were detected or not detected in the sediments. Amongst ASVs from the surface water column, the relative abundance of taxa at subdivision level are presented for those that were B) detected in the sediments (represented by dark blue in A) and C) not detected in the sediments (represented by light blue in A). Subtropical and subantarctic water masses were analysed separately.

Within the shared ASVs common between upper water column, fixed traps and sediments (i.e., protists from the upper water column exported via sinking particles to the sediments), the community composition was different between sample types (Figure 6B). The relative abundance values reported below correspond to the subsets presented in Figures 6B and C. Haptophyta and Gyrista had similar relative abundance in fixed trap and sediment samples between the water masses. Haptophyta (mainly Prymnesiophyceae) decreased in relative abundance from fixed traps to the sediments (31% and 0.4%, respectively). In contrast, Gyrista (mainly Mediophyceae) increased in relative abundance from fixed traps to the sediments (6.8% and 19.5%, respectively). Mediophyceae in the sediments were dominated by *Minidiscus variabilis* (Thalassiosirales) in SA-Sc, and *Chaetoceros dichatoensis* (Chaetocerotales) in all other cycles. Green algae and other heterotrophic groups had several differences in the community composition between water masses. For phototrophic groups, fixed traps and sediment samples in subtropical cycles had a low relative abundance of Mamiellophyceae (*<*5%). In subantarctic cycles, fixed traps had a higher proportion of reads from Chloropicophyceae (mainly *Chloroparvula pacifica*; 12.9%) and Prasinodermophyceae (only *Prasinoderma singularis*; 3.2%), which increased by at least 4-fold in sediment samples (45.2% and 23.7%, respectively). For heterotrophic groups, fixed traps in subtropical cycles were dominated by Radiolaria (mainly Polycystinea, Acantharia and RAD-A at 31.4%, 10.1% and 6.1%, respectively), while sediment samples had a lower proportion of reads from Radiolaria, except in ST2 with 40% of reads from Polycystinea (mainly *Cladococcus scoparius*), and a higher proportion of reads from Labyrinthulomycetes (Bigyra) at 43.9%. In subantarctic cycles, fixed traps were dominated by Ciliophora (41.8%) which decreased by 23-fold in sediment samples (1.8%).

Within shared ASVs common only between the upper water column and fixed traps (i.e., upper water column protists which were lost during export to the sediments), Ciliophora dominated in nearly all cycles of fixed trap samples (average relative abundance of 58.5%; Figure 6C). Radiolaria had a higher proportion of reads in subantarctic compared to subtropical cycles (average of 38.3% and 5.4%, respectively), while the relative abundance of Gyrista peaked in SA-Sc in fixed traps at 34.5%, composed of *Pseudo-nitzschia* sp. (Bacillariophyceae), unclassified Thalassiosirales sp. (Mediophyceae), and unclassified Hemidiscaceae sp. (Coscinodiscophyceae).

Amongst ASVs common between the lower water column and fixed traps (i.e., protists from the lower water column exported via sinking particles), Radiolaria (Polycystinea and Acantharia) and Ciliophora dominated the community, which together accounted for 98.6% of reads and 83% of ASVs (Figures S7 and Table S2). Considering the total protist community in fixed trap samples (including Dinoflagellata), Ciliophora ASVs from the upper water column had a higher relative abundance in fixed trap samples compared to ASVs from the lower water column (11.3% and 3.0%, respectively), but the opposite was observed for Radiolaria ASVs (9.2% and 14.6%, respectively).

Similar to the rest of the protist community, Dinoflagellata (Dinophyceae and Syndiniales) ASVs from the upper water column had 6-fold higher relative abundance and twice the number of ASVs in the fixed traps and sediment samples compared to the lower water column (Figure S8A). In the fixed trap samples, shared Dinophyceae ASVs (from upper and lower water column) had a 4-fold higher proportion of reads compared to Syndiniales (Figure S8B-D), but half the number of ASVs (Table S2). In the sediments, shared Syndiniales ASVs instead had a 6-fold higher proportion of reads and twice the number of ASVs compared to Dinophyceae. This suggests most of the Dinophyceae exported from the upper and lower water column were lost during export to the sediments, whereas Syndiniales with a low proportion in fixed trap samples had accumulated in the sediments.

### Changes in ASV relative abundance during export and transfer

We used upper water column and fixed trap samples to further investigate the changes in relative abundance of particle-associated protist taxa from the surface to mesopelagic depths. We applied differential abundance (DESeq2) analysis separately for each water mass (subtropical and subantarctic) to obtain the predicted change in relative abundance (log2-fold change; LFC) of ASVs between 1) the mixed layer of the upper water column and upper fixed traps (below the mixed layer at depth levels 1 and 2; refer to Table 1) and 2) the upper fixed traps and lower fixed traps (upper mesopelagic, depth levels 3 and 4), to infer export and transfer potential of taxa at ASV level (Figures S9, S10, S11 and S12). ASVs with higher and lower relative abundance in the upper water column compared to upper fixed traps were characterised as “low” and “high export”, respectively. ASVs with higher and lower relative abundance in the upper fixed traps compared to the lower fixed traps were characterised as “low” and “high transfer”, respectively. As the community was dominated by Dinoflagellata across all sample types, Dinoflagellata ASVs were presented separately in Figure 7C-D.

**Figure 7:**
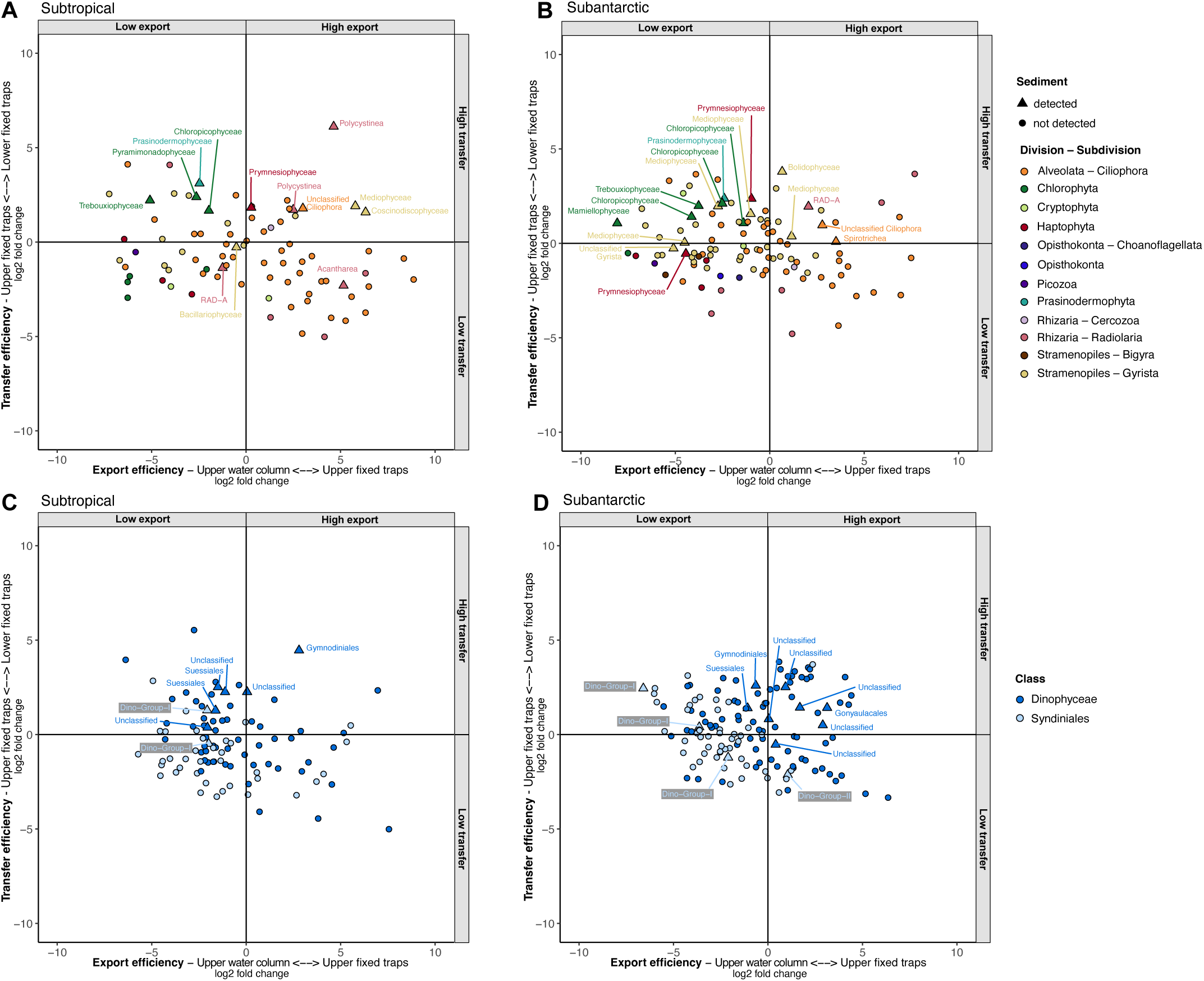
Predicted differential abundance analysis (DESeq2) of protist ASVs between the upper water column (5 to 40 m) and upper fixed traps (70 m to 100 m) on the x-axis, and between the upper fixed traps and lower fixed traps (300 m to 500 m) on the y-axis, for protist ASVs excluding Dinoflagellata in A) subtropical and B) subantarctic cycles, and for Dinoflagellata ASVs in C) subtropical and D) subantarctic cycles. The colours represent the subdivision or class level of the ASVs, and the shapes represent whether the ASV was detected in the sediments of the same water mass. DeSEQ2 for each analysis are presented in Figures S9, S10, S11 and S12.

We obtained 187 and 245 ASVs from the analysis for subtropical and subantarctic cycles respectively, of which 103 ASVs were shared between the two water masses (Table S3). The majority of ASVs were assigned to Dinoflagellata (180, 100 ASVs from Dinophyceae and 80 ASVs from Syndiniales) and Ciliophora (63, 54 ASVs from Spirotrichea), with a lower number of ASVs from Gyrista (42), Radiolaria (15), Chlorophyta (10) and Haptophyta (7).

Nearly all the taxa had similar patterns of change in predicted differential abundance for export (upper water column vs. upper fixed traps) and transfer (upper fixed traps vs. lower fixed traps) in subtropical and subantarctic water masses (Figure 7 and Table S4). The mean LFC of Phaeocystales (Haptophyta) and Syndiniales ASVs showed low export and low transfer. The mean LFC of Gyrista, Prasinodermophyta (1 ASV, present in both subtropical and subantarctic) and Chloropicophyceae (Chlorophyta) ASVs had low export and high transfer. The mean LFC of Ciliophora and Radiolaria ASVs showed high export and low transfer. Dinophyceae ASVs were dominated by Gymnodiniales and unclassified Dinophyceae, which showed no clear difference in LFC for export and transfer (Figure S11 and S12). Mamiellophyceae (Chlorophyta) and Isochrysidales (Haptophyta) showed different patterns between the water masses based on mean LFC values. Mamiellophyceae ASVs had low export in both subtropical and subantarctic, but low transfer in subtropical and high transfer in subantarctic. Isochrysidales ASV (assigned to the same *Gephyrocapsa huxleyi* ASV) had high transfer in both subtropical and subantarctic, but high export in subtropical and low export in subantarctic.

Fourty-one out of 329 ASVs obtained from the analysis were detected in sediment samples (Figure 7). They belonged to Dinoflagellata (12 ASVs to Dinophyceae and 5 ASVs to Syndiniales), Chlorophyta (6), Radiolaria (5) and Gyrista (8), with a lower number of ASVs assigned to Ciliophora (2), Haptophyta (2) and Prasinodermophyta (1). About 80% of these ASVs were characterised as high transfer.

### Influence of remineralisation on particle-associated 18S rRNA genes

Live traps were the only sample type where Metazoans (66.9%) dominated the community over protist taxa, with Arthropoda representing 50.4% of the total eukaryotic reads. We compared the relative abundance and ASV richness of the protist community between fixed and live traps to identify the effects of particle degradation and consumption. Among ASVs common between both fixed and live traps, ASV richness decreased by 45.7*±*28% from fixed to live traps. Dinophyceae and Syndiniales (Dinoflagellata) had a slightly higher than average loss of ASV richness (57.5% and 61.6%, respectively), but also gained 78 and 157 ASVs unique to live traps (Table S5). Radiolaria and Ciliophora had a 3 and 10-fold loss in relative abundance, and 54.6% and 76.5% loss of ASV richness (Table S5), respectively.

Among photosynthetic taxa, Isochrysidales, Mediophyceae and Thalassiosirales had similar relative abundance in live and fixed traps, while Chloropicophyceae, Prasinodermophyceae and Chrysophyceae represented a higher proportion of reads in live compared to fixed traps (Figures 4D, 5). Nearly all Mamiellophyceae, Chloropicophyceae and Prasinodermophyceae ASVs from fixed traps were found in live traps, while approximately half of diatom ASVs from fixed traps were found in live traps (Bacillariophyceae and Mediophyceae at 55.6% and 31.4%, respectively; Table S5).

Bicoecea (Anoecales) and Phaeodarea (Phaeodendrida) were two heterotrophic orders that had higher proportion of reads and ASV richness in live traps (Figure 5 and Table S5). Phaeodaria (*Coelodendrum ramosissimum*) was only detected in high proportion in ST1 (22.9%), while the abundance of Anoecales (*Caecitellus paraparvulus*) was only high in SA2 and low for the other cycles (0.04-1%). There was also a low proportion of reads assigned to Labyrinthulomycetes across all cycles (0.02-0.4%).

## Discussion

### High impact of upper water column protist community on export

Our results indicate that the particle-associated protist diversity exported to the sediments was accumulated from the suspended protist community at the surface mixed layer, with lesser influence from the lower water column community at mesopelagic and bathypelagic depths where they sink through (Figure 6). This trend contrasts with patterns reported for bacteria in Li et al. (2023), which showed across a 3-year time series that the particle-associated bacterial community from bathypelagic depths was dominated by suspended bacteria from mesopelagic to bathypelagic depths over those from upper water column, also similarly suggested in LeCleir et al. (2014) and Stephens et al. (2024). Suspended microbial communities from the lower water column can accumulate in sinking particles through colonisation and growth influenced by grazing (Artolozaga et al. 1997; Bochdansky et al. 2017; Nguyen et al. 2022; Pernice et al. 2015), as well as forming new or repackaging sinking particles through particle aggregation and metazoan faecal pellet production (Huffard et al. 2020; Wilson et al. 2013). The succession of particle-associated bacterial communities during export indicates a preference toward bacteria with slow growth rates and anaerobic metabolisms, suited to the conditions in bathypelagic depths and more degraded particles (Boeuf et al. 2019; Stephens et al. 2024). Based on the lower water column taxa detected in sinking particles which are mixo- and phagotrophic (Dinophyceae, Radiolaria, and Ciliophora), a possible reason for this difference is that these taxa prefer to feed on fragmented suspended particles over sinking particles (“microbial gardening”, see Mayor et al. (2014)). It is also possible that these taxa have physical characteristics or specific interactions which could have led to loss through detachment, grazing from metazoans or mortality.

While a high proportion of upper water column taxa were exported in sinking particles across the mixed layer, most of the protist diversity (90% of ASVs) were lost during transfer from below the mixed layer to the sediments. The eukaryotic community that was exported to the sediments was likely heavily repackaged during their downward transfer (Cordier et al. 2022; Yang et al. 2024), as the community composition of shared ASVs in the upper water column, sinking particles and sediments were different between the three sample types (Figure 6B). The relative abundance of protists in sinking particles were dominated by phago-mixotrophic and phagotrophic groups (Dinophyceae, Radiolaria, and Ciliophora), which have all been described to dominate the community of sinking particles in previous sediment trap studies (Boeuf et al. 2019; Duret et al. 2020; Durkin et al. 2022; Lampitt et al. 2023; Poff et al. 2021; Preston et al. 2020; Valencia et al. 2022; Yang et al. 2024). The exported protist taxa in the sediments were instead represented by phytoplankton groups of green microalgae (Chlorophyta and Prasinodermophyta) and diatoms (Mediophyceae), as well as parasitic Syndiniales, which have been detected in sinking particles and in the sediments, mostly at lower proportion compared to the overall community especially for phytoplankton groups (Cordier et al. 2022; Durkin et al. 2022; Preston et al. 2020; Valencia et al. 2021). Our results highlight the benefits of jointly surveying the sinking particles via sediment traps and deep ocean sediments, as contents of the sediment traps might not be representative of the protist taxa which contributes to deep ocean carbon export.

Considering this change in the exported community between sinking particles and those deposited in the sediments, we further explored how the protist community transformed during export from surface mixed layer to mesopelagic depths. The change in relative abundance of eukaryotic assemblages during export showed different patterns of accumulation between taxa at different depths (Figure 7), which could imply unequal loss of taxa through remineralisation, particle fragmentation (Duret et al. 2020), and/or selective consumption (Gutierrez-Rodriguez et al. 2019). As particles of different sizes can have a distinct community composition likely controlled by food web interactions (Durkin et al. 2022), it could also affect the export potential of taxa, as well as the sinking speeds and microbial interactions within the particle during transfer (Bach et al. 2019; Guidi et al. 2009; Weber et al. 2016). Surprisingly, these export and transfer patterns were similar within each taxon in contrasting water masses, regardless of the surface relative abundance and export flux. The previous studies mentioned which reported unequal taxa loss though particle fragmentation and selective consumption (Duret et al. 2020; Gutierrez-Rodriguez et al. 2019), as well as distinct community composition in different particle sizes (Durkin et al. 2022), also observed such similar community patterns in productive and oligotrophic areas, including HNLC conditions, indicating that the drivers for protist loss during export are conserved across environmental gradients. Furthermore, this and previous studies also showed sample type had a greater influence on protist communities compared to water masses with contrasting biogeochemical properties and POC fluxes (Fontanez et al. 2015; Gutierrez-Rodriguez et al. 2019; Valencia et al. 2021), overall indicating that the community which influences export can be similar in different water masses.

### Taxonomic diversity of exported communities across the water masses Phytoplankton

#### Diatoms

The contribution of diatoms toward export flux is well established (Duret et al. 2020; Durkin et al. 2022; Nelson et al. 1995; Poff et al. 2021; Preston et al. 2020; Yang et al. 2024) and is largely influenced by their large cell size, silica-rich skeleton and bloom forming capabilities (Durkin et al. 2021). Diatom-enriched particles are also suggested to have more efficient export with less particle fragmentation (Duret et al. 2020), and rapid export of active or resting stage cells (Rodríguez-Martínez et al. 2020). In this study, the diatom community followed the same pattern between the upper water column and fixed traps, with a high proportion of large pennate diatoms (Bacillariales), large centric diatoms (Coscinodiscophyceae) and small to mid-sized centric diatoms (Mediophyceae), suggesting no preference in export of any diatom taxon over others. However, there were differences in the diatom community exported to the sediments compared to the community lost during export. ASVs assigned to *Pseudo-nitzschia* sp. (Bacillariophyceae), unclassified Thalassiosirales (Mediophyceae), and unclassified Hemidiscaceae (Coscinodiscophyceae), mostly belonging to larger diatoms, were not detected in the sediments, whereas the small nano-diatom *Minidiscus variabilis* (Thalassiosirales) and the chain-forming *Chaetoceros dichatoensis* (Chaetocerotales) were present in the sediments. Both *Minidiscus* sp. and *Chaetoceros* sp. are globally correlated to POC export to the sediments (Cordier et al. 2022). *Chaetoceros dichatoensis* was not present or present in very low abundance in the upper water column and fixed trap samples, suggesting that it was exported to the sediment before sampling had occurred and stored in the sediment as resting spores (Wilks et al. 2021). *Minidiscus variabilis* was enriched in the upper water column and fixed trap samples of SA-Sc, showing continuous export of the diatom to the sediments in this cycle. The export of small diatoms, including *Minidiscus*, are probably due to their incorporation into large particles through faecal pellets or aggregation (Durkin et al. 2022; Smayda 1971). In contrast, larger diatoms such as *Pseudo-nitzschia* sp. (Bacillariophyceae) and Coscinodiscophyceae were not found in the sediment of this study, despite dominating the same upper water column samples as *Minidiscus* with almost 3-fold higher relative abundance. This suggests that while a large diatom size can boost the sinking of cells (Goldman 1993), it does not necessarily translate into export to the sediments (Leblanc et al. 2018; Lin et al. 2017; Williams et al. 2024), which could be due to its enrichment in smaller particles over larger particles (Durkin et al. 2022) or contradictory effects from diatom buoyancy regulation (Williams et al. 2024).

#### Green algae

Chloropicophyceae and Prasinodermophyceae present in the surface water column of subantarctic cycles were abundantly exported to subantarctic sediments, while Mamiellophyceae that were dominant in surface water column samples of subtropical cycles were present in the sediments at very low abundance. In particular, the export of Prasinodermophyceae appears to be disproportionately higher (35-fold) compared to its surface abundance, in comparison to other small phytoplankton taxa that also accumulated in the sediments (Chloropicophyceae and *Minidiscus variabilis*, 3-fold increase). The aggregation of small phytoplankton to larger particles, has been strongly suggested as a mechanism for small phytoplankton to be exported out of the surface water community (Lomas and Moran 2011; Richardson and Jackson 2007), occurring year-round (Li et al. 2023). The same small phytoplankton taxa have also been detected in low proportion in sinking particles in previous studies (Preston et al. 2020; Valencia et al. 2022, 2021), in particular in the larger particles (Durkin et al. 2022). However, this would not explain the differences in the deposition of Chloropicophyceae and Prasinodermophyceae in the sediment compared to Mamiellophyceae, especially since all three taxonomic groups have similar loss of ASV richness in live traps compared to fixed traps, suggesting similar rates of degradation. The relative abundance of Chloropicophyceae in fixed trap samples was likely underestimated as they were not properly fixed with formalin, as the relative abundance of Chloropicophyceae was higher in lower water column and live trap samples (Figure 4).

#### Haptophyta

Haptophyta species *Gephyrocapsa huxleyi* and *Phaeocystis antarctica* are prevalent phytoplankton genera in the region (Chang and Northcote 2016; Gutiérrez-Rodríguez et al. 2022) and commonly detected in sinking particles (Arrigo et al. 1999; Duret et al. 2020; Gowing et al. 2001; Rigual Hernández et al. 2018). These species had different export patters, where *Phaeocystis antarctica* had “low transfer” patterns, while *Gephyrocapsa huxleyi* had “high transfer” patterns (Figures S9 and S10). Although extensive blooms of *Phaeocystis antarctica* can result in rapid export of this taxon via aggregates to mesopelagic depths (Asper and Smith Jr. 1999; DiTullio et al. 2000), under non-blooming conditions, *Phaeocystis*-enriched particles seem to be subjected to higher occurrences of particle fragmentation (Duret et al. 2020; Yang et al. 2024) and hence not likely to be exported to the sediment. On the other hand, *Gephyrocapsa huxleyi* is ballasted by its CaCO_3_ coccoliths, which increase sinking speed (Buitenhuis et al. 2001) and therefore reduces remineralisation pressure (Martin et al. 1987), leading to increased enrichment in sinking particles at deeper depths (Yang et al. 2024). Moreover, the high export of *Gephyrocapsa huxleyi* across the mixed layer in subtropical compared to subantarctic waters was likely boosted by the presence of salps in cycle ST2 (see Décima et al. 2023). Despite this difference, both *Gephyrocapsa huxleyi* and *Phaeocystis antarctica* were found in very low proportion in the sediments. Microscopy observations in the same region have also reported low contributions of cocolithopore to export flux at lower mesopelagic depths (Wilks et al. 2021).

### Protozooplankton

The relative abundance of Dinophyceae, Radiolaria and Ciliophora were higher in fixed traps compared to upper water column samples, suggesting that these taxa were preferentially exported out of the upper water column (Figure 7). Dinophyceae and Radiolaria were constantly identified in sinking particles at abyssal depths across a 9-month sampling period, during which the surface communities and export fluxes changed, suggesting constant contribution of these taxa to export (Preston et al. 2020). Despite overwhelmingly dominating fixed trap samples, surface ASVs from Dinophyceae, Radiolaria and Ciliophora were detected at less than half the proportion of photosynthetic taxa in the sediments. This suggests Dinophyceae, Radiolaria and Ciliophora as important components of sinking particles, possibly transforming the community within the particles and as a source of organic carbon to mesopelagic communities, but does not consistently contribute to the sequestration of organic carbon in the sea floor in this region especially when compared to photosynthetic taxa.

#### Radiolaria

The high relative abundance of Radiolaria in sinking particles agrees with previous observations (Amacher et al. 2009; Duret et al. 2020; Fontanez et al. 2015; Gutierrez-Rodriguez et al. 2019; Lampitt et al. 2009; Preston et al. 2020; Valencia et al. 2022). As Radiolaria usually have a higher relative abundance in the lower compared to the upper water column across the open ocean (Flegontova et al. 2023; Giner et al. 2020; Obiol et al. 2020; Ollison et al. 2021), including at this sampling location (Gutiérrez-Rodríguez et al. 2022), this taxa is suggested to have a lower than perceived export contribution as they are underrepresented in the fixed traps that coincide with their depth distribution maxima (Valencia et al. 2021). Indeed, we find that there was a slightly higher proportion of reads from lower compared to upper water column Radiolarian ASVs, suggesting the entire water column as an important source for the particle-associated Radiolarian community. Despite the high proportion of Radiolaria reads in sinking particles, the relative abundance of Radiolaria was overall very low in the sediment. This was similarly observed in Preston et al. (2020), which reported that periods where Radiolaria dominated in sinking particles did not correspond with increasing POC flux nor aggregation on the sea floor. Based on network analysis, Polycystinea and Acantharia are suggested to be disproportionately affected by parasitism from Syndiniales during export, and therefore have higher remineralisation and loss compared to other protist hosts (Anderson et al. 2024). The exception was RAD-A, which was the only group consistently exported throughout the cycles and present in the sediments throughout all cycles, although at a lower proportion. RAD-A was also noted as having consistent presence in sinking particles (Gutierrez-Rodriguez et al. 2019) across seasons (Preston et al. 2020), with low associations with Syndiniales during export (Anderson et al. 2024), which could suggest low but consistent export of RAD-A across seasons and biogeochemically different water masses.

#### Ciliophora

Ciliophora in sinking particles were dominated by Oligotrichia and Choreotrichida, both of which contain species which are either phago-mixotrophic or heterotrophic (Schneider et al. 2020), and OLIGO5 which are mainly heterotrophic. Ciliates are usually enriched in sinking particles (Duret et al. 2020; Gowing et al. 2001), likely feeding on bacteria and small protists (Caron et al. 2012), thus playing an important role in trophic transfer and nutrient recycling in the dark ocean. Surprisingly, the lower relative abundance of Ciliophora ASVs in mesopelagic compared to mixed layer fixed traps, as well as the high proportion of ASV richness lost in live traps, suggests loss of ciliates during particle sinking. In contrast, ciliate richness and abundance tends to increase in live traps compared to fixed traps (Gutierrez-Rodriguez et al. 2019; Valencia et al. 2021), as ciliates are known to have a dominant role in the consumption of primary biomass and the production of sinking particles (Zeldis and Décima 2020). Although Ciliophora are usually reported to be abundant in the lower water column (Grattepanche et al. 2016; Gutiérrez-Rodríguez et al. 2023; Ollison et al. 2021), they are in very low abundance in the lower water column samples of this study (Table S2). The loss of Ciliophora during particle sinking could be due to selective and rapid consumption by copepods (Zeldis and Décima 2020), as copepod ASVs were enriched in the live trap samples.

#### Dinoflagellata

Dinoflagellata (Dinophyceae and Syndiniales) are distributed throughout the water column (Cohen et al. 2021; Flegontova et al. 2023; Obiol et al. 2020) and overwhelmingly dominated all sample types of this study in both water masses (Figures 4 and 5). The ecological functions of the two taxa are very different, as Dinophyceae have phototrophic, mixotrophic and phagotrophic trophic modes (Schneider et al. 2020), while Syndiniales are parasites to a wide range of hosts, including Radiolaria, diatoms, Dinophyceae and ciliates (Berdjeb et al. 2018; Bråte et al. 2012; Harada et al. 2007; Sehein et al. 2022). Dinophyceae and Syndiniales are suggested to have important ecological roles in carbon export, as they are dominant in the sediment traps deployed across seasons and contrasting environments (Duret et al. 2020; Durkin et al. 2022; Preston et al. 2020; Valencia et al. 2021; Yang et al. 2024). While Dinophyceae has positive correlation with organic carbon export, the correlation of parasitic Syndiniales is less certain (Anderson et al. 2024; Guidi et al. 2016), indicating the contribution of Syndiniales toward carbon flux is via both export and remineralisation.

In this study, Syndiniales had “low export” patterns and lower relative abundance in fixed trap samples compared to Dinophyceae, which suggests a lower contribution toward export of this parasitic group, although they can have the same relative abundance as Dinophyceae in sinking particles (Duret et al. 2020; Poff et al. 2021; Preston et al. 2020). Euphotic Dinophyceae engage in both phototrophy and phagotrophy, while mesopelagic Dinophyceae show an upregulation of phagotrophic behaviours (Cohen et al. 2021; Jeong et al. 2010). As Dinophyceae can alter their nutrient metabolism to adapt to different environmental conditions (Cohen et al. 2021), this could suggest a shift of surface Dinophyceae from phototrophy to phagotrophy as particles are sinking, affecting the remineralisation depth of the particle. In the sediments, particle-associated Syndiniales dominated in terms of ASV richness and proportion of reads over Dinophyceae instead. Similarly, Cordier et al. (2022) found a higher proportion of parasitic protists ASVs that were sinking from the surface compared to those that were not sinking, overall indicating a disproportionate diversity of parasitic protists being exported to the sediments.

### Eukaryotic community transformation during particle degradation

Sinking particles are areas of elevated microbial and metazoan activity (Karl and Knauer 1984; Taylor et al. 1986), where live traps in this study showed an enrichment of heterotrophic, parasitic and saprotrophic protists, as well as zooplankton, consistent with the transformations of sinking particles reported in previous studies (Fontanez et al. 2015; Gutierrez-Rodriguez et al. 2019). Live trap samples were substantially enriched with copepod ASVs, which are known to ingest and fragment sinking organic particles (Lampitt et al. 1990) and therefore strongly influence carbon flux attenuation (Bressac et al. 2024). The fragmentation of sinking organic particles by copepods could be a strategy to increase the abundance of particle-associated bacteria, flagellates, and ciliates for later grazing in a strategy known as microbial gardening (Mayor et al. 2014; Zeldis and Décima 2020), as observed with the increase in relative abundance of the heterotrophic nanoflagellate species *Caecitellus paraparvulus* and of other small flagellate groups (Filosa). There was also an increase in the ASV richness and relative abundance of Phaeodaria (*Coelodendrum ramosissimum*), which likely feed on small flagellates enriched around the sinking particles (Gowing and Bentham 1994). Syndiniales were also more enriched in live traps with new ASVs not detected in fixed traps, possibly due to the introduction of other hosts. Lastly, there was a higher proportion of reads and ASVs of Labyrinthulomycetes in live traps compared to fixed traps, suggesting an increase in saprotrophic activity originating from the lower water column, contributing to the degradation of organic matter in sinking particles (Bochdansky et al. 2017). In subtropical cycles, an upper water column ASV assigned to the Labyrinthulomycetes genus *Aplanochytrium* which is widely distributed in pelagic waters (Bai et al. 2022), was also detected in sinking particles (at a very low proportion of 0.015%) and sediments (3.5%; Figure 6B). Labyrinthulomycetes are known to be active and significant contributors to the upper layers of deep ocean sediments (Rodríguez-Martínez et al. 2020). This could indicate a unique role for Labyrinthulomycetes as both contributors to vertical export and remineralisers of sinking organic material (Bai et al. 2022), similar to Syndiniales.

The protist community between fixed and live sediment traps were significantly different as observed previously (Fontanez et al. 2015; Gutierrez-Rodriguez et al. 2019; Valencia et al. 2021), suggesting that the microbial community on sinking particles is modified during export. While Ciliophora, Radiolaria, and diatoms (especially Mediophyceae) had greater loss of ASV richness, suggesting these groups had more rapid remineralisation, dominant green photosynthetic groups, including Chloropicophyceae, Mamiellophyceae and Prasinodermophyceae, did not appear to be influenced by degradation. This was also observed by Durkin et al. (2022), who compared ASV richness of particles that were incubated on the ship. The loss in ASV richness between live and fixed traps, and also between fixed traps and sediment samples, is assumed to be a reflection of the organic material lost from the taxa. However, it is possible that DNA of an organism is degraded but its organic material remains (see discussion in Valencia et al. 2021).

In conclusion, our results highlight three key takeaways. Firstly, in contrast to bacteria (Li et al. 2023), a 2-fold higher relative abundance in the particle-associated eukaryotic community was attributed to surface water column ASVs compared to lower water column ASVs, implying that the suspended protist community from the lower water column minimally influenced particle-associated eukaryotic community. Secondly, most of the diversity was lost during particle sinking from below the mixed layer to the sediment. However, the loss of diversity was not similar across taxonomic groups. Despite the high relative abundance of phago-mixotrophic and heterotrophic taxa (Dinophyceae, Radiolaria and Ciliophora) in fixed trap samples (as previously reported in Boeuf et al. 2019; Duret et al. 2020; Gutierrez-Rodriguez et al. 2019; Lampitt et al. 2009, 2023; Poff et al. 2021; Valencia et al. 2022), particle-exported taxa in the sediments were dominated by pico and nano photosynthetic taxa of Mediophyceae (except for *Chaetoceros dichatoensis*), Chloropicophyceae and Prasinodermophyceae. Interestingly, surface ASVs assigned to parasitic Syndiniales and the saprotrophic Labyrinthulomycetes genus *Aplanochytrium* had higher relative abundance in sediments compared to fixed trap samples, indicating that these heterotrophic taxa might have been accumulated in the sediments from surface export (Bai et al. 2022; Cordier et al. 2022). Lastly, export and transfer patterns differed between taxa, suggesting unequal loss of taxa through remineralisation, selective consumption (Gutierrez-Rodriguez et al. 2019) or particle fragmentation (Duret et al. 2020) during export. This pattern was similar within each taxonomic group in both oligotrophic subtropical and HNLC subantarctic water masses regardless of the relative abundance in surface, which further emphasizes that the drivers for protist loss during export are not strongly related to environmental conditions or surface community, but possibly to specific microbial interactions, the type of particle exported in and/or inherent cell properties that are conserved across environmental gradients.

## Acknowledgements

We acknowledge the crew of RV *Tangaroa* for their efforts during the TAN1810 voyage. We thank Daniel Vaulot (CNRS, France) for feedback on data processing. We are grateful to Debbie Hulston (NIWA, New Zealand) for sample processing support.

DRYO and ALS were supported by RG91/21 and RG26/19 Singapore Ministry of Education, Academic Research Fund Tier 1. AGR was supported by NIWA via the New Zealand Ministry of Business, Innovation and Employment’s Strategic Science Investment Funding to the National Coasts Ocean Centre.

## Availability of data and materials

Source code (all sequence processing and analysis scripts) and raw data produced for this paper is available on Github (https://github.com/deniseong/TAN1810_sedtraps). All unprocessed sequencing data have been deposited in NCBI Sequence Read Archive (PRJNA670061 and PRJNA1033349).

## Author contributions statement

ALS, AGR, SN have contributed to conception and study design. ALS, AGR, JB, SN, MRS, MD have processed the samples and produced data. DRYO, AGR, ALS, SN, MRS, MD have analysed and interpreted the data. DRYO drafted the manuscript. All authors have revised and approved the final manuscript.

