## Supplementary Material for "Export dynamics of protists across the southern subtropical frontal zone reveal taxon-specific patterns"

#### Materials and methods

##### Particle Interceptor Traps

The PITs were deployed once at the start of each experimental cycle (only in SA-Sc-a of cycle SA-Sc) to collect sinking particles for DNA analysis and assess export fluxes of particulate organic carbon (POC), Chlorophyll *a* (Chl *a*) and phaeopigment concentration. The PIT arrays were deployed for 2.8 to 4 days at four depths (Table 2), typically starting from 30 m below the mixed layer depth (estimated by CTD profiles of temperature, salinity and fluorescence, 70 m for almost all the cycles), as well as 100 m, 300 m and 500 m below the surface (Table 1). The PIT arrays consisted of 12 PIT cylindrical traps (inner diameter: 7 cm and length: 58 cm) on each cross-frame, with one cross-frame for each depth. Each cylindrical trap was baffled by smaller tubes (internal diameter: 1.3 cm, length: 7.5 cm) at the top of the cylinder. Prior to deployment, each cylinder was filled with a high density brine solution to just below the bottom of the baffles, with or without formalin (0.4% formaldehyde final concentration) for fixed (DNA, POC, Chl *a* and phaeopigment) and live (DNA) traps, respectively. The brine solution was prepared by combining surface seawater passed through a SupraPak 0.2 $\mu$m filter with 50 g $L^{-1}$ NaCl solids to +5‰ above ambient density. Additionally, borax was added as a buffer to a final concentration of 1 g $L^{-1}$ to reduce acidification of carbonates, and $SrSO_4$ was added to a final concentration of 72 mg $L^{-1}$ to minimise the dissolution of Acantharia. Upon recovery, each trap was processed separately. The overlying seawater of traps were aspirated down to within 2 cm of the top of the brine, leaving a total sample volume of 600-700 mL per trap. The contents were filtered though a 200 $\mu$m mesh and the meshes were viewed under a dissecting microscope (×20 magnification) to manually remove zooplankton “swimmers”. The remaining material on the meshes were recombined with the filtered trap contents and filtered on pre-combusted 25 mm GF/F filters for POC analysis (1 full trap each), uncombusted 25 mm GF/F filters for Chl *a* and phaeopigments (50 mL subsample), 0.2 $\mu$m 142 mm polycarbonate membrane filter for total contents of fixed and live traps (1 full trap each) and polyethersulphone 47 mm filters with 20 and 0.2 $\mu$m pore size sequentially for size-fractionated fixed and live traps (half trap each). Samples for Chl *a*, phaeopigments and DNA were stored at -80 °C until processing, and POC samples were processed and its export fluxes calculated as described in Décima et al. (2023). There were no live trap samples collected for SA-Sc.

##### Multicorer

Sediment samples were collected with an Ocean Instruments MC-800 multicorer once each cycle (only in SA-Sc-a of cycle SA-Sc) (Table 1), where 6 core tubes of 10 cm diameter were deployed. Before sediment sampling, seawater was aspirated off each core to a height of 1 cm above the sediment surface. After which, the sample for DNA sequencing was obtained from one core, where a surface scrape removed 25 mL of sediment into a 50 mL Falcon tube and stored at -80 °C until processing.

##### Amplicon analysis

Sequences were processed on RStudio Version 1.4.1717 (RStudio Team 2021). Primer sequences were removed from raw sequences using *Cutadapt* Version 3.4 (Martin 2011). Fastq files were trimmed and quality filtered using the function 'filterandtrim' with the *DADA2* R package Version 1.12 (Callahan et al. 2016). Reads were trimmed and filtered to the following options: truncLen = c(220, 210), minLen = c(220, 210), truncQ = 2, maxEE = c(10, 10). Forward and reverse sequences were then dereplicated, grouped and merged with default settings. Chimeras were identified and removed using the function 'removeBimeraDenovo', resulting in 8,656 ASVs. Taxonomy was assigned with the function 'assignTaxonomy' against the PR2 database version 5.0 (Guillou et al. 2013). ASVs with a bootstrap value of $<$ 90% at supergroup level were removed, resulting in 7,400 ASVs. ASVs with low bootstrap support ($<$ 80%) at the species level were reclassified to a higher taxonomic level where the bootstrap values were higher or equals to 80%. ASVs with less than 10 reads were removed from this study. ASVs assigned to Domain Bacteria (3 ASVs) and subdivision Fungi (96 ASVs) were not considered. ASVs assigned to *Gephyrocapsa oceanica* with 100% bootstrap support at species level were reassigned as *Gephyrocapsa huxleyi* (previously known as *Emiliana huxleyi*, Bendif et al. 2019) since *G. oceanica* and *G. huxleyi* are indistinguishable using the 18S rRNA gene. *G. huxleyi* was chosen mainly based on microscopy observations (Chang and Northcote 2016) and higher number of cultures of this species isolated in the region (Gutiérrez-Rodríguez et al. 2022). We excluded fixed trap samples from SA2 at 540 m depth as they were dominated by a single ASV and had a much lower diversity than the rest of the fixed trap samples. Analysis was performed with *Phyloseq* R package Version 1.36.0 (McMurdie and Holmes 2013) and *microViz* Version 0.10.10 (Barnett et al. 2021).

Formalin-fixed and preservative-free sediment traps were processed via two filtration methods to obtain the total ($>$ 0.2µm) and size fractionated (pico-nano: 0.2-20 µm and micro: $>$ 20µm) community. Within fixed traps, the community composition between total and micro samples were significantly different (PERMANOVA-Adonis, $R^2$ = 0.05, P = 0.03). Pico-nano size fraction was not considered in this analysis because of the low number of samples (n = 4) relative to micro (n = 19) and total (n = 13), however the community pico-nano samples were more similar to micro compared to total (Figure S4 and S1A-C). The main difference in the community composition between the samples was the high proportion of Radiolaria in the total community (49.8±26.3%), a common feature across most of the total community samples collected (Figure S4), which was surprisingly in low proportion in both pico-nano (7.1±4.9%) and micro (15.8±10.7%) size fractionated samples. Gutiérrez-Rodríguez et al. (2019) had recovered high proportion of Radiolaria sequences in both size fractionated community (< 0.8 and > 0.8 µm) of fixed traps, where they obtained each fraction independently, compared to this study where we used sequential filtration instead. The size-fractionating methodology applied in this study likely influenced in particular the Radiolaria community within fixed traps. Live trap samples had no significant differences in the protist community between the three size-fractions (Figure S1D-F) (PERMANOVA-Adonis, $R^2$ = 0.04, P = 0.78). We have averaged the community within fixed and live traps across all size fractions for all analysis.

#### Methodological considerations

With the extensive sampling conducted, the comparisons between sample types depicted in this study should also be carefully interpreted due to the differences in 1) time and space between the sample types, and 2) sampling methods. DNA reads in the sediment could have been deposited up to 40 days prior before sampling (see discussion in Décima et al. 2023), and possibly for thousands of years depending on the environmental conditions (Torti et al. 2015), accumulating DNA from organisms not present in the upper water column and sediment traps during the time of sampling. However, as the sinking speeds of particles can differ by more than 200 times (1 to 200 m day$^{-1}$, Asper 1987; Asper and Smith Jr. 1999). Therefore, there can be a disconnection between the fixed trap communities sampled at different depths, for example in ST2 where the sediment traps at 300 m likely contained some particles which were not originating from the surface waters of the same water mass (Figure 2). We only analysed the presence and absence of ASVs, but not their abundance, between the surface and lower water column sampled with Niskin and high-volume McLane pumps, respectively, due to the differences in volume of water collected. The microbial community obtained with CTD compared to high-volume McLane pumps is shown to capture different types of particles, with in situ pumps capturing a higher diversity and micro-sized particles (Mackinson et al. 2015; Puigcorbé et al. 2020). Despite the differences in sampling methodology, the high relative abundance of Radiolaria, Dinophyceae and Syndiniales from McLane pump samples showed similar patterns to that of mesopelagic depths sampled with Niskin bottles (Flegontova et al. 2023; Giner et al. 2020; Obiol et al. 2020). Despite these limitations, the combination of these different sampling strategies in this study should provide useful insight into the succession of the different plankton groups in during particle sinking, and the various sources of its community.

##### Eukaryotic composition of the water column and sediments

In the upper water column, Dinoflagellata dominated the community (56.0±12.2%), with Dinophyceae (35.4±9.7%) at slightly higher proportion than Syndiniales (20.3±11.1%), followed by Gyrista (11.1±6.5%), Chlorophyta (10.8±8.8%), Haptophyta (9.5±4.5%) and Ciliophora (5.4±2.7%; Figure 4A). The relative abundance of Dinoflagellata and Ciliophora were fairly consistent across cycles, while photosynthetic taxa (Chlorophyta, Gyrista and Haptophyta) varied (Figure 5). Chlorophyta were dominated by picoplanktonic groups (Mamiellophyceae and Chloropicophyceae). Mamiellophyceae (*Micromonas commoda* A2, *Ostreococcus* sp. and *Bathycoccus prasinos*) were more abundant in subtropical compared to subantarctic cycles and peaked in SA-Sc at 22%, while Chloropicophyceae (mainly *Chloroparvula pacifica*) had higher abundance in SA1 and SA2. Diatoms consisted mostly of Bacillariophyceae (*Pseudo-nitzschia* sp. and *Cylindrotheca closterium*), which had the highest relative abundances cycles SA-Sc and SA1 (8-11%), and Mediophyceae (*Minidiscus variabilis*) with higher relative abundance only in SA-Sc (3%). Within Haptophyta, Isochrysidales (only *Gephyrocapsa huxleyi*) and Prymnesiales (mainly *Chrysochromulina* sp.) were more abundant in subtropical cycles, while Phaeocystales (mainly *Phaeocystis antarctica*) were more abundant in subantarctic cycles.

In the lower water column, Dinoflagellata (59.7±22%) and Radiolaria (30.8±25.5%) overwhelmingly dominated the protist community across all the cycles (Figure 4B). The proportion of reads of Syndiniales (42.8±18.5%) was higher than Dinophyceae (16.5±12%). Within Radiolaria, Polycystinea (15.2±20.9%) and RAD-A (11.2±20.7%) were the major taxa, with minor contributions from Acantharia (2.1±2.3%). Ciliophora had very low abundance (¡ 1%). Within the photosynthetic community, only Chloropicophyceae (mainly *Chloroparvula pacifica*) was present at high relative abundance of up to 25% in SA1 from depths 70 m to 300 m (Figures 5 and S4B).

In the sediment samples, Cercozoa (40.9±9.7%) and Dinoflagellata (21.6±6.2%) dominated the community (Figure 4E). The majority of reads from Cercozoa were from Filosa-Thecofilosea (21.7±8.7%) and Filosa-Imbricatea (12.5±15.2%), nearly all of which were unique to sediment samples. Labyrinthulomycetes were present across all cycles (2-7%). Within Dinoflagellata, Syndiniales (17±7%) had 3-fold higher abundance compared to Dinophyceae (4.5±2.6%). Ciliophora were present across all the cycles (4.2-11%) and consisted of Scuticociliatia, which were abundant only in sediment samples, and Spirotrichea

(Figure 5). There was a low proportion of reads assigned to Radiolaria across all cycles (0.65-7.3%), mostly from Polycystinea and RAD-A. Photosynthetic taxa present included Chloropicophyceae, Prasinodermophyceae, Chrysophyceae, and diatoms (Mediophyceae and Bacillariophyceae). Chloropicophyceae (mainly *Chloroparvula pacifica*) and Prasinodermophyceae (only *Prasinoderma singularis*) had the highest proportion of reads from photosynthetic taxa, which were mostly present in subantarctic cycles (4-11% and 3-7%, respectively). Within Mediophyceae, the proportion of Thalassiosirales (mostly *Minidiscus variabilis*) peaked in SA-Sc at 7%, while Chaetocerotales (mostly *Chaetoceros dichatoensis*) abundance was more consistent across the cycles (0.6-3%). Chrysophyceae (0.07-0.6% across all cycles) and Bacillariophyceae (0.1-0.2% only in ST2 and SA-Sc) had a lower proportion of reads.

### Tables

**Table S1:** Depths (m) sampled in the surface water column for various measurements at each cycle.

| Cycle | Nutrient concentration | Size-fractionated Chlorophyll *a* | Net primary production (standard volume) | Filtered samples for DNA analysis |
|---|---|---|---|---|
| | Depth (m) | Depth (m) | Depth (m) | Depth (m) |
| ST1 | 5, 10, 12, 20, 25, 30, 35, 40, 50, 60, 100, 160, 200 | 10, 20, 25, 30, 40, 50, 70, 100 | 5, 12, 20, 25, 30, 35, 40, 50 | 5, 12, 20, 25, 30, 35, 40 |
| ST2 | 5, 10, 12, 20, 25, 30, 40, 50, 70, 100 | 10, 25, 40, 50, 70, 100 | 5, 12, 20, 25, 30, 40, 50 | 5, 12, 20, 30, 35, 40 |
| SA-Sc | 5, 10, 12, 20, 25, 30, 40, 50, 55, 70, 100, 200, 500 | 10, 25, 40, 50, 70, 100 | 5, 10, 12, 20, 30, 40, 50 | 5, 12, 20, 30, 35, 40 |
| SA1 | 5, 10, 12, 20, 25, 30, 40, 50, 60, 70, 80, 100 | 10, 25, 40, 50, 70, 100 | 5, 12, 20, 25, 30, 40, 60 | 5, 12, 20, 30, 40 |
| SA2 | 5, 10, 12, 20, 25, 45, 50, 70, 90, 100, 160, 200 | 10, 25, 50, 70, 80, 100, 160, 200 | 5, 12, 25, 30, 45, 50, 60, 70, 90, 100 | 5, 12, 25, 30 |

**Table S2:** Number of protistan ASVs in **fixed trap samples** that are shared between upper (sampled with Niskin) and lower (sampled with McLane pump) water column samples, or not detected in the water column samples (unique). ASVs detected in water column samples are further divided into ASVs that are detected or not detected in the sediments.

| | | Upper water col. | | Lower water col. | | | |
|---|---|---|---|---|---|---|---|
| Subdivision | Class | Detected in sediments | Not detected in sediments | Detected in sediments | Not detected in sediments | Unique | Total |
| Alveolata_X | Alveolata_XX | 0 | 0 | 0 | 0 | 3 | 3 |
| Alveolata_X | Ellobiopsidae | 0 | 1 | 0 | 0 | 0 | 1 |
| Ancyromonadida_X | Ancyromonadida_XX | 0 | 0 | 0 | 0 | 3 | 3 |
| Apicomplexa | Apicomplexa_X | 0 | 1 | 0 | 0 | 0 | 1 |
| Apusomonada_X | Apusomonadidae | 0 | 1 | 0 | 0 | 0 | 1 |
| Archaeplastida_XX | Archaeplastida_XXX | 0 | 1 | 0 | 0 | 0 | 1 |
| Bigyra | Sagenista | 1 | 0 | 3 | 3 | 2 | 9 |
| Bigyra | Opalozoa | 0 | 5 | 0 | 0 | 0 | 5 |
| Bigyra | Bicoecea | 1 | 1 | 0 | 0 | 2 | 4 |
| Centroplasthelida_X | Pterocystida | 0 | 1 | 0 | 0 | 0 | 1 |
| Cercozoa | Filosa-Thecofilosea | 1 | 2 | 1 | 1 | 1 | 6 |
| Cercozoa | Phaeodarea | 0 | 0 | 0 | 3 | 3 | 6 |
| Cercozoa | Cercozoa_X | 0 | 3 | 0 | 1 | 1 | 5 |
| Cercozoa | Filosa-Granofilosea | 1 | 1 | 0 | 0 | 0 | 2 |
| Cercozoa | Filosa-Imbricatea | 0 | 0 | 1 | 0 | 0 | 1 |
| Chlorophyta_X | Mamiellophyceae | 3 | 4 | 0 | 0 | 0 | 7 |
| Chlorophyta_X | Chloropicophyceae | 5 | 0 | 0 | 1 | 0 | 6 |
| Chlorophyta_X | Pyramimonadophyceae | 1 | 3 | 0 | 0 | 1 | 5 |
| Chlorophyta_X | Chlorophyta_XX | 0 | 1 | 0 | 0 | 0 | 1 |
| Chlorophyta_X | Prasino-Clade-VIII | 0 | 1 | 0 | 0 | 0 | 1 |
| Chlorophyta_X | Trebouxiophyceae | 1 | 0 | 0 | 0 | 0 | 1 |
| Choanoflagellata | Choanoflagellatea | 0 | 5 | 0 | 0 | 1 | 6 |
| Ciliophora | Spirotrichea | 1 | 107 | 1 | 22 | 157 | 288 |
| Ciliophora | Oligohymenophorea | 0 | 11 | 0 | 12 | 92 | 115 |
| Ciliophora | Phyllopharyngea | 1 | 14 | 0 | 0 | 21 | 36 |
| Ciliophora | CONThreeP | 0 | 9 | 0 | 1 | 13 | 23 |
| Ciliophora | Ciliophora_X | 1 | 5 | 0 | 0 | 6 | 12 |
| Ciliophora | Litostomatea | 0 | 3 | 0 | 0 | 2 | 5 |
| Ciliophora | CONTH_8 | 0 | 2 | 0 | 0 | 1 | 3 |
| Ciliophora | Heterotrichea | 0 | 2 | 0 | 1 | 0 | 3 |
| Ciliophora | Prostomatea | 0 | 2 | 0 | 0 | 0 | 2 |
| Ciliophora | Cyclotrichium_like_organism | 0 | 0 | 0 | 0 | 1 | 1 |
| Cryptophyta_X | Cryptophyceae | 0 | 5 | 0 | 0 | 2 | 7 |
| Dinoflagellata | Syndiniales | 21 | 255 | 26 | 143 | 90 | 535 |
| Dinoflagellata | Dinophyceae | 14 | 197 | 3 | 49 | 106 | 369 |
| Dinoflagellata | Dinoflagellata_X | 0 | 3 | 0 | 8 | 2 | 13 |
| Dinoflagellata | Noctilucophyceae | 0 | 4 | 0 | 0 | 1 | 5 |
| Gyrista | Mediophyceae | 6 | 22 | 2 | 2 | 2 | 34 |
| Gyrista | Bacillariophyceae | 1 | 13 | 0 | 1 | 2 | 17 |
| Gyrista | Coscinodiscophyceae | 2 | 8 | 2 | 2 | 0 | 14 |

**Table S2:** *(continued)*

| | | Upper water col. | | Lower water col. | | | |
|---|---|---|---|---|---|---|---|
| | | Detected in sediments | Not detected in sediments | Detected in sediments | Not detected in sediments | | |
| Subdivision | Class | | | | | Unique | Total |
| Gyrista | Gyrista_X | 1 | 11 | 1 | 0 | 1 | 14 |
| Gyrista | Chrysophyceae | 0 | 6 | 1 | 0 | 3 | 10 |
| Gyrista | Pelagophyceae | 0 | 4 | 0 | 0 | 1 | 5 |
| Gyrista | Dictyochophyceae | 0 | 3 | 0 | 0 | 1 | 4 |
| Gyrista | Bolidophyceae | 1 | 1 | 0 | 0 | 1 | 3 |
| Gyrista | MOCH-2 | 0 | 2 | 0 | 0 | 0 | 2 |
| Gyrista | Peronosporomycetes | 0 | 1 | 0 | 0 | 0 | 1 |
| Gyrista | MOCH-3 | 1 | 0 | 0 | 0 | 0 | 1 |
| Gyrista | Pirsoniales | 0 | 0 | 0 | 0 | 1 | 1 |
| Haptophyta_X | Prymnesiophyceae | 2 | 8 | 0 | 1 | 0 | 11 |
| Haptophyta_X | Haptophyta_Clade_HAP3 | 0 | 1 | 0 | 0 | 0 | 1 |
| Kathablepharida | Kathablepharidea | 0 | 2 | 0 | 1 | 0 | 3 |
| Opisthokonta_X | Opisthokonta_XX | 0 | 1 | 0 | 3 | 8 | 12 |
| Picozoa_X | Picozoa_XX | 0 | 4 | 0 | 0 | 0 | 4 |
| Prasinodermophyta_X | Prasinodermophyceae | 1 | 0 | 0 | 0 | 0 | 1 |
| Radiolaria | Acantharea | 2 | 20 | 2 | 33 | 44 | 101 |
| Radiolaria | Polycystinea | 4 | 6 | 11 | 45 | 25 | 91 |
| Radiolaria | RAD-A | 2 | 3 | 0 | 21 | 1 | 27 |
| Radiolaria | RAD-B | 0 | 2 | 0 | 12 | 0 | 14 |
| Radiolaria | RAD-C | 0 | 1 | 1 | 4 | 0 | 6 |
| Radiolaria | Radiolaria_X | 0 | 0 | 0 | 3 | 0 | 3 |
| Streptophyta_X | Embryophyceae | 0 | 0 | 0 | 1 | 0 | 1 |
| TSAR_XX | TSAR_XXX | 0 | 0 | 0 | 2 | 0 | 2 |
| Telonemia_X | Telonemia_XX | 0 | 5 | 0 | 2 | 2 | 9 |
| Tubulinea_X | Elardia | 0 | 0 | 0 | 0 | 5 | 5 |
| | Total | 75 | 774 | 55 | 378 | 608 | 1890 |

**Table S3:** Number of ASVs obtained from DESeq2 analysis for each water mass grouped by subdivision and class.

| Subdivision | Class | ST | SA | Both | Total |
|---|---|---|---|---|---|
| Bigyra | Opalozoa | 0 | 2 | 0 | 2 |
| Cercozoa | Unclassified | 0 | 0 | 1 | 1 |
| Chlorophyta | Mamiellophyceae | 1 | 1 | 2 | 4 |
| Chlorophyta | Chloropicophyceae | 0 | 2 | 1 | 3 |
| Chlorophyta | Pyramimonadophyceae | 2 | 0 | 0 | 2 |
| Chlorophyta | Trebouxiophyceae | 0 | 0 | 1 | 1 |
| Choanoflagellata | Choanoflagellatea | 0 | 1 | 0 | 1 |
| Ciliophora | Spirotrichea | 18 | 15 | 21 | 54 |
| Ciliophora | CONThreeP | 2 | 1 | 1 | 4 |
| Ciliophora | Phyllopharyngea | 0 | 1 | 0 | 1 |
| Ciliophora | Oligohymenophorea | 1 | 0 | 0 | 1 |
| Ciliophora | Prostomatea | 1 | 0 | 0 | 1 |
| Ciliophora | Unclassified | 0 | 0 | 1 | 1 |
| Ciliophora | Litostomatea | 0 | 0 | 1 | 1 |
| Cryptophyta | Cryptophyceae | 1 | 1 | 1 | 3 |
| Dinoflagellata | Dinophyceae | 17 | 38 | 45 | 100 |
| Dinoflagellata | Syndiniales | 27 | 42 | 11 | 80 |
| Dinoflagellata | Noctilucophyceae | 1 | 0 | 0 | 1 |
| Gyrista | Mediophyceae | 1 | 8 | 5 | 14 |
| Gyrista | Bacillariophyceae | 0 | 10 | 2 | 12 |
| Gyrista | Coscinodiscophyceae | 0 | 5 | 2 | 7 |
| Gyrista | Unclassified | 1 | 3 | 0 | 4 |
| Gyrista | Bolidophyceae | 0 | 1 | 0 | 1 |
| Gyrista | Dictyochophyceae | 0 | 1 | 0 | 1 |
| Gyrista | MOCH-2 | 1 | 0 | 0 | 1 |
| Gyrista | Pelagophyceae | 1 | 0 | 0 | 1 |
| Gyrista | Chrysophyceae | 0 | 0 | 1 | 1 |
| Haptophyta | Prymnesiophyceae | 2 | 2 | 2 | 6 |
| Haptophyta | Haptophyta_Clade_HAP3 | 0 | 1 | 0 | 1 |
| Opisthokonta | Unclassified | 0 | 1 | 0 | 1 |
| Picozoa | Unclassified | 0 | 0 | 1 | 1 |
| Prasinodermophyta | Prasinodermophyceae | 0 | 0 | 1 | 1 |
| Radiolaria | Acantharea | 3 | 4 | 0 | 7 |
| Radiolaria | Polycystinea | 3 | 2 | 0 | 5 |
| Radiolaria | RAD-A | 0 | 0 | 2 | 2 |
| Radiolaria | RAD-B | 0 | 0 | 1 | 1 |
| | Total | 83 | 142 | 103 | 328 |

**Table S4:** Log2fold change values (mean±SD) obtained from DESeq analysis for the main taxa.

| | | | Subtropical | | | Subantarctic | | |
|---|---|---|---|---|---|---|---|---|
| Subdivision | Class | Order | Export LFC | Transfer LFC | No. of ASVs | Export LFC | Transfer LFC | No. of ASVs |
| Chlorophyta | Chloropicophyceae | | -1.97 | 1.67 | 1 | -2.68 ± 1.38 | 1.53 ± 0.53 | 3 |
| Chlorophyta | Mamiellophyceae | | -6.23 ± 0.06 | -2.29 ± 0.59 | 3 | -8.57 ± 1.38 | 0.38 ± 0.81 | 3 |
| Chlorophyta | Trebouxiophyceae | | -5.09 | 2.21 | 1 | -3.77 | 1.99 | 1 |
| Ciliophora | | | 1.48 ± 3.43 | -0.88 ± 1.93 | 46 | 0.81 ± 3.31 | -0.27 ± 1.74 | 41 |
| Dinoflagellata | Dinophyceae | | -0.41 ± 2.71 | 0.04 ± 2.01 | 62 | 0.13 ± 2.61 | 0.64 ± 1.89 | 83 |
| Dinoflagellata | Syndiniales | | -1.56 ± 2.79 | -1.18 ± 1.42 | 38 | -2.53 ± 1.94 | -0.11 ± 1.59 | 53 |
| Gyrista | | | -1.71 ± 4.19 | 0.64 ± 1.46 | 14 | -2.08 ± 2.65 | 0.34 ± 1.34 | 38 |
| Haptophyta | Prymnesiophyceae | Isochrysidales | 0.27 | 1.82 | 1 | -0.97 | 2.37 | 1 |
| Haptophyta | Prymnesiophyceae | Phaeocystales | -4.42 | -2.03 | 1 | -3.8 ± 0.57 | -1.26 ± 0.95 | 3 |
| Prasinodermophyta | | | -2.47 | 3.11 | 1 | -2.41 | 2.37 | 1 |
| Radiolaria | | | 3.46 ± 4.7 | -0.67 ± 3.83 | 9 | 3.82 ± 5.48 | -1.78 ± 4.07 | 9 |

**Table S5:** Change in ASV richness between fixed and live traps.

| Subdivision | Class | Proportion of fixed trap ASVs in live traps | Total no. of ASVs in fixed traps | No. of ASVs unique to live traps |
|---|---|---|---|---|
| Ancyromonadida_X | Ancyromonadida_XX | 100 | 3 | 1 |
| Apusomonada_X | Apusomonadidae | 100 | 1 | 4 |
| Bigyra | Bicoecea | 100 | 4 | 5 |
| Bigyra | Sagenista | 44.4 | 9 | 21 |
| Bigyra | Opalozoa | 40 | 5 | 7 |
| Centroplasthelida_X | Pterocystida | 100 | 1 | 0 |
| Cercozoa | Filosa-Granofilosea | 100 | 2 | 3 |
| Cercozoa | Filosa-Imbricatea | 100 | 1 | 5 |
| Cercozoa | Phaeodarea | 83.3 | 6 | 10 |
| Cercozoa | Cercozoa_X | 80 | 5 | 2 |
| Cercozoa | Filosa-Thecofilosea | 33.3 | 6 | 12 |
| Chlorophyta_X | Chloropicophyceae | 100 | 6 | 4 |
| Chlorophyta_X | Trebouxiophyceae | 100 | 1 | 0 |
| Chlorophyta_X | Mamiellophyceae | 85.7 | 7 | 1 |
| Chlorophyta_X | Pyramimonadophyceae | 40 | 5 | 3 |
| Choanoflagellata | Choanoflagellatea | 16.7 | 6 | 1 |
| Ciliophora | Litostomatea | 40 | 5 | 0 |
| Ciliophora | Phyllopharyngea | 34.2 | 38 | 9 |
| Ciliophora | CONThreeP | 21.7 | 23 | 2 |
| Ciliophora | Ciliophora_X | 16.7 | 12 | 1 |
| Ciliophora | Oligohymenophorea | 15.7 | 115 | 2 |
| Ciliophora | Spirotrichea | 12.4 | 290 | 8 |
| Cryptophyta_X | Cryptophyceae | 42.9 | 7 | 1 |
| Dinoflagellata | Dinophyceae | 42.5 | 374 | 78 |
| Dinoflagellata | Syndiniales | 38.4 | 544 | 157 |
| Dinoflagellata | Dinoflagellata_X | 30.8 | 13 | 4 |
| Dinoflagellata | Noctilucophyceae | 20 | 5 | 0 |
| Gyrista | MOCH-3 | 100 | 1 | 0 |
| Gyrista | Pirsoniales | 100 | 1 | 2 |
| Gyrista | Pelagophyceae | 60 | 5 | 3 |
| Gyrista | Bacillariophyceae | 55.6 | 18 | 1 |
| Gyrista | Coscinodiscophyceae | 53.3 | 15 | 0 |
| Gyrista | Chrysophyceae | 50 | 10 | 4 |
| Gyrista | Bolidophyceae | 33.3 | 3 | 1 |
| Gyrista | Mediophyceae | 31.4 | 35 | 2 |
| Gyrista | Dictyochophyceae | 20 | 5 | 2 |
| Gyrista | Gyrista_X | 20 | 15 | 1 |
| Haptophyta_X | Prymnesiophyceae | 36.4 | 11 | 6 |
| Kathablepharida | Kathablepharidea | 33.3 | 3 | 0 |
| Opisthokonta_X | Opisthokonta_XX | 33.3 | 12 | 1 |
| Picozoa_X | Picozoa_XX | 25 | 4 | 2 |
| Prasinodermophyta_X | Prasinodermophyceae | 100 | 1 | 1 |
| Radiolaria | RAD-C | 66.7 | 6 | 1 |
| Radiolaria | RAD-B | 64.3 | 14 | 1 |
| Radiolaria | RAD-A | 40.7 | 27 | 9 |
| Radiolaria | Acantharea | 34.7 | 101 | 8 |
| Radiolaria | Radiolaria_X | 33.3 | 3 | 1 |
| Radiolaria | Polycystinea | 32.3 | 93 | 12 |
| Streptophyta_X | Embryophyceae | 100 | 2 | 0 |
| TSAR_XX | TSAR_XXX | 50 | 2 | 1 |
| Telonemia_X | Telonemia_XX | 11.1 | 9 | 1 |
| Tubulinea_X | Elardia | 100 | 5 | 9 |

### Figures

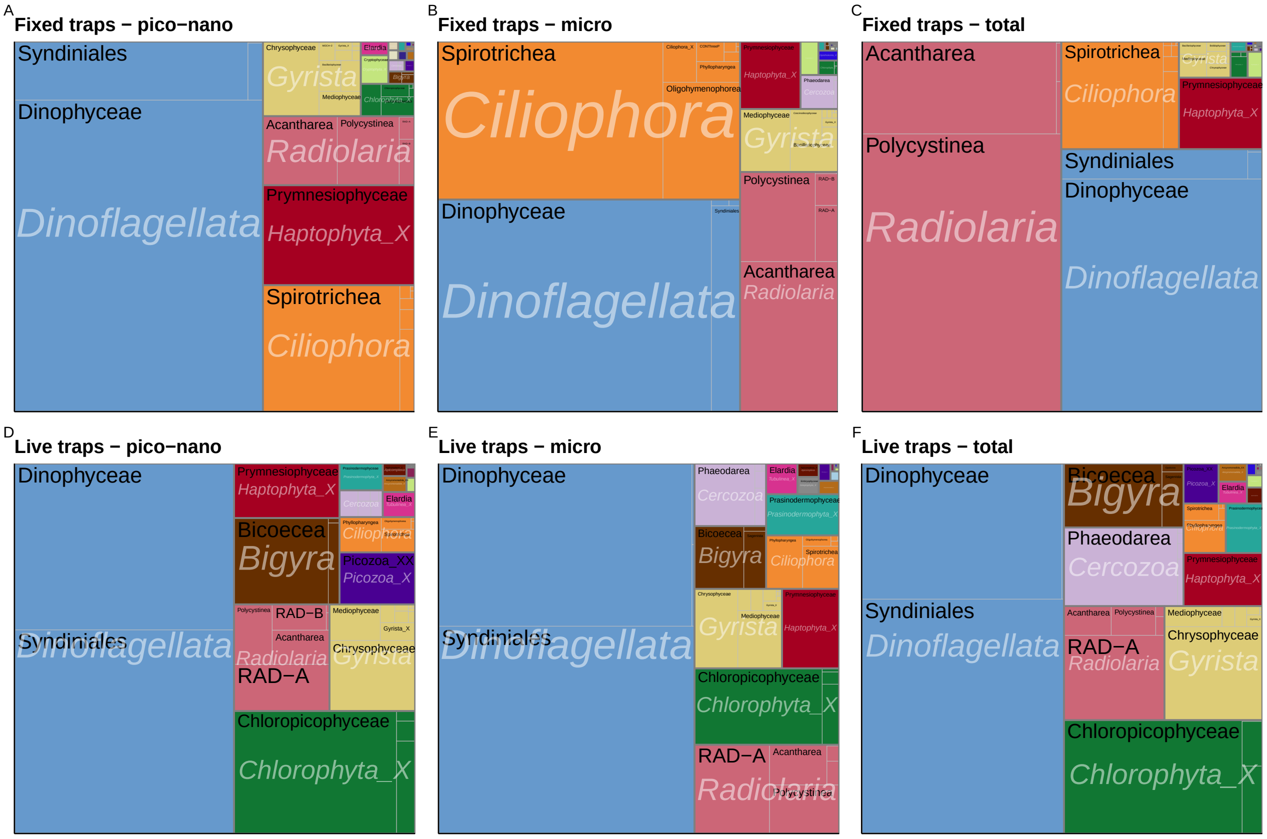


**Figure S1:** Treemap of 18S V4 rRNA protist community at division (text in white italics) and class (black text) level for each size fraction (pico-nano, micro and total) from fixed (A, B, C) and live (D, E, F) traps samples. Rectangles within each treemap are proportional to the relative abundance of each taxa.

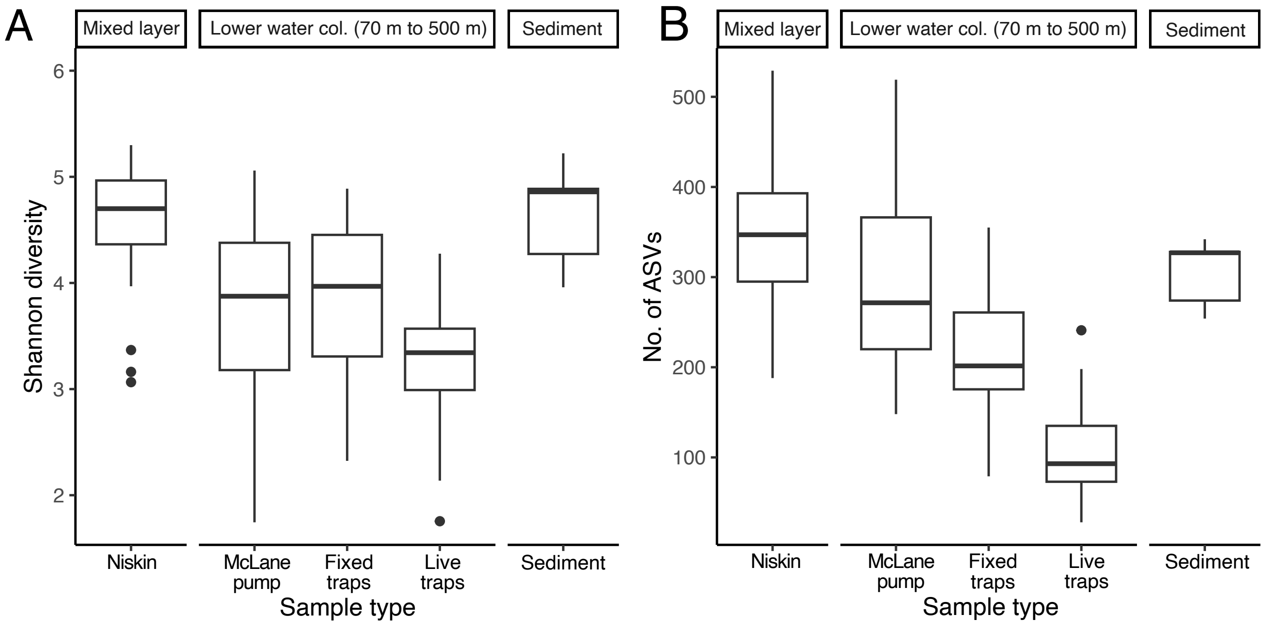


**Figure S2:** (A) Shannon diversity and (B) richness of samples based on protist ASVs as a function of sample type and grouped by depth sampled.

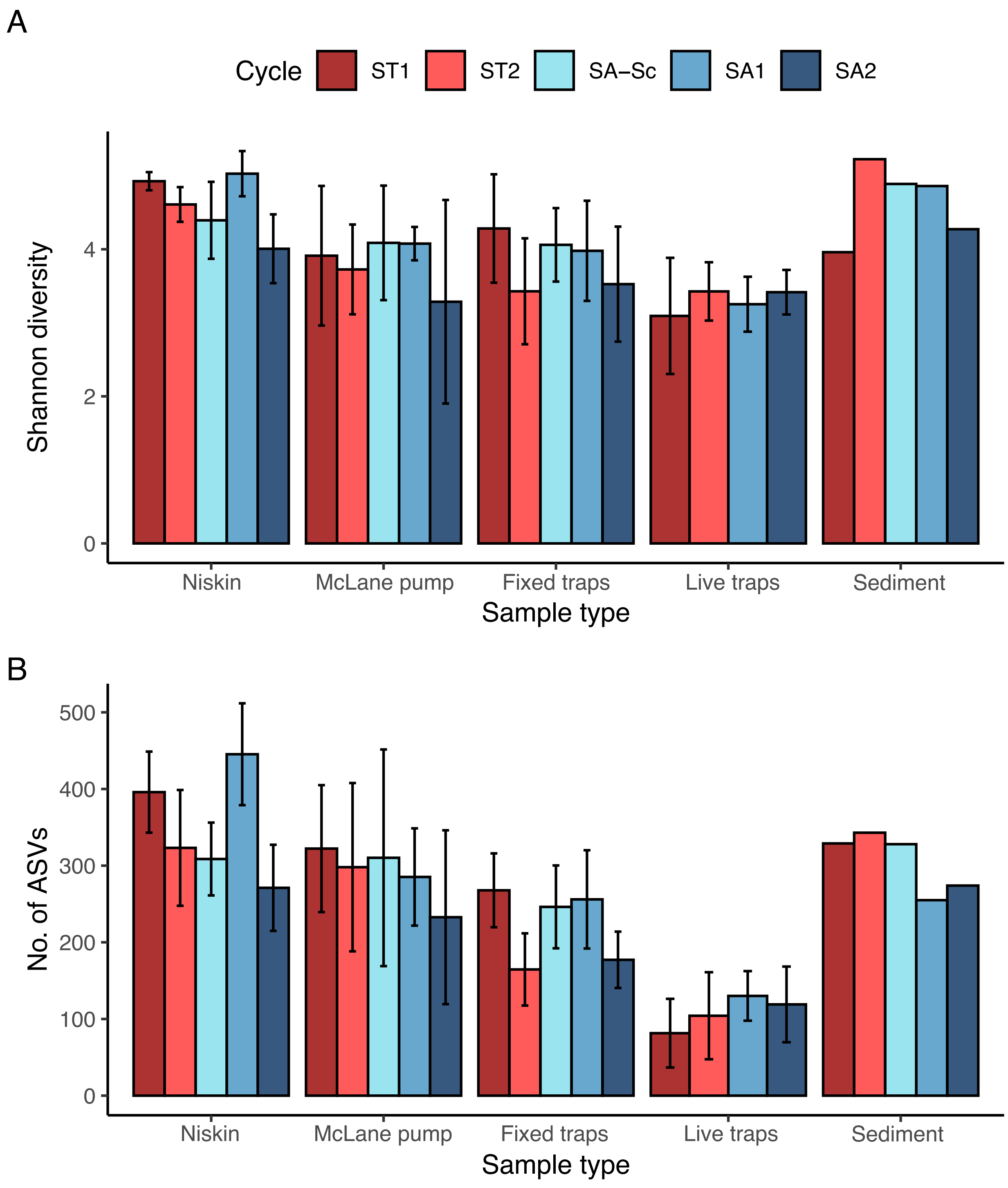


**Figure S3:** (A) Shannon diversity and (B) number of ASVs of samples in each cycle (indicated by colour) and sample type.

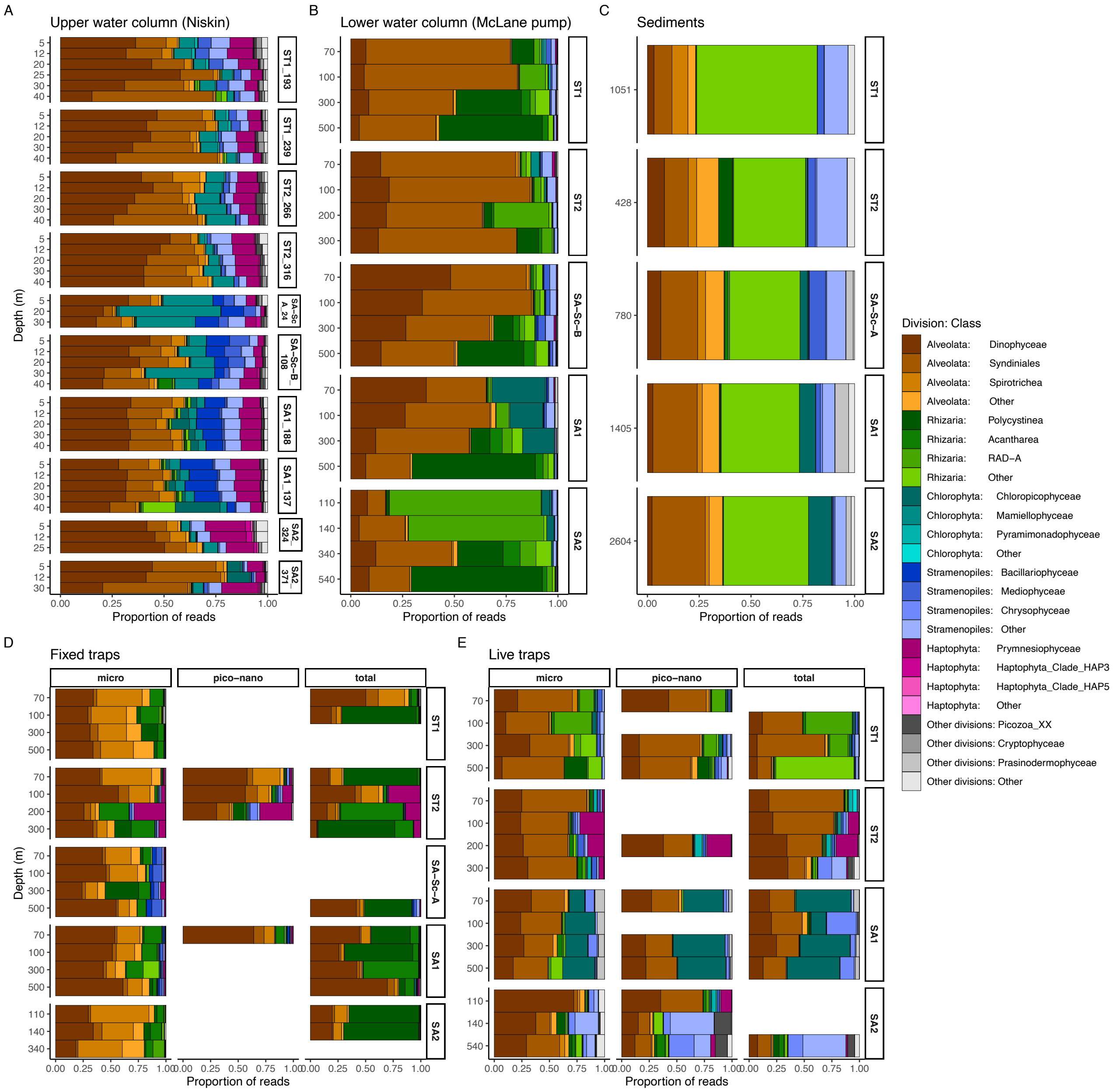


**Figure S4:** Relative abundance of 18S V4 rRNA protist community of samples from each sample type and cycle. Taxa is presented at class level and its respective colours are grouped at division level.

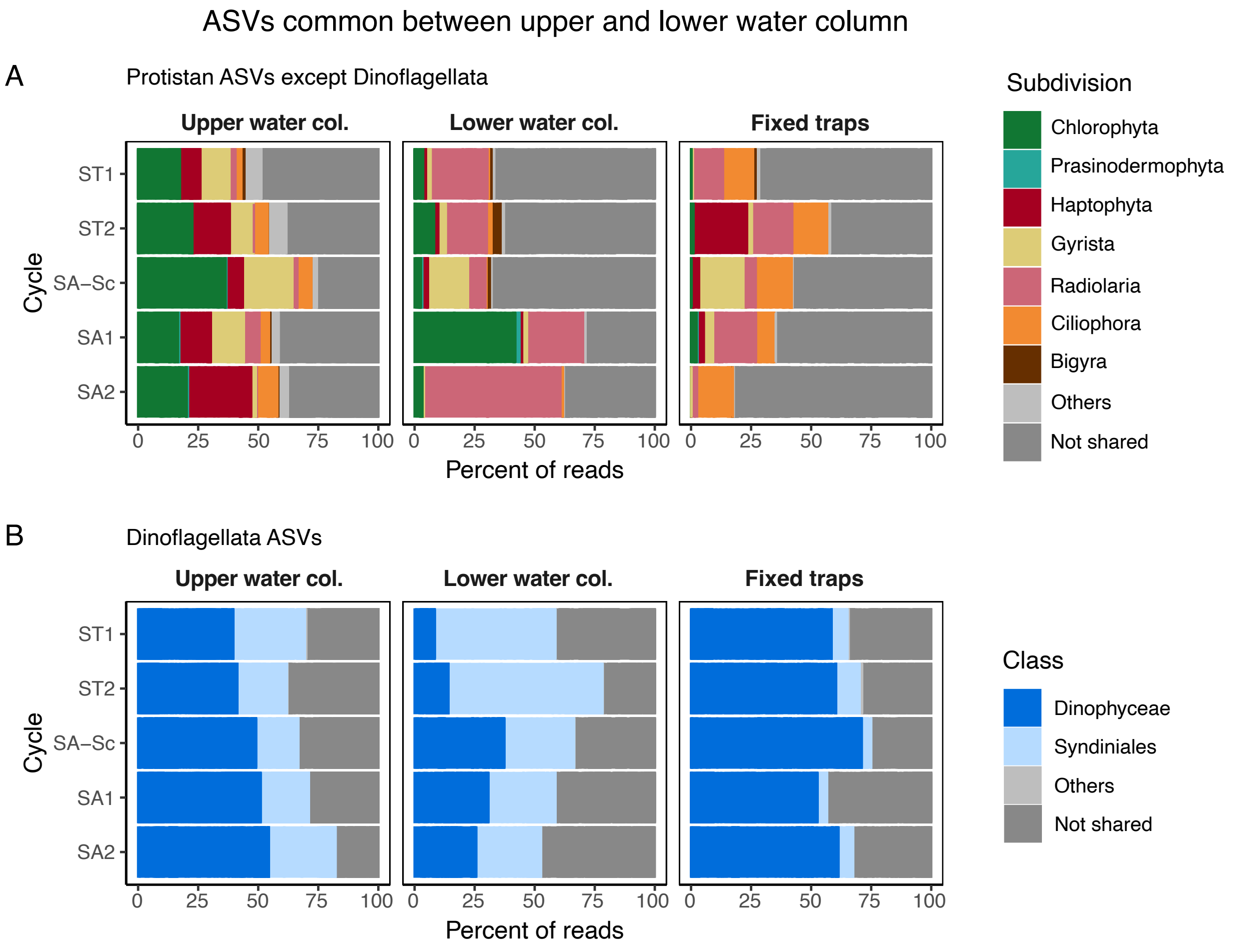


**Figure S5:** The relative abundance of (A) protist ASVs excluding Dinoflagellata and (B) Dinoflagellata ASVs, shared between the upper water column, lower water column and fixed trap samples.

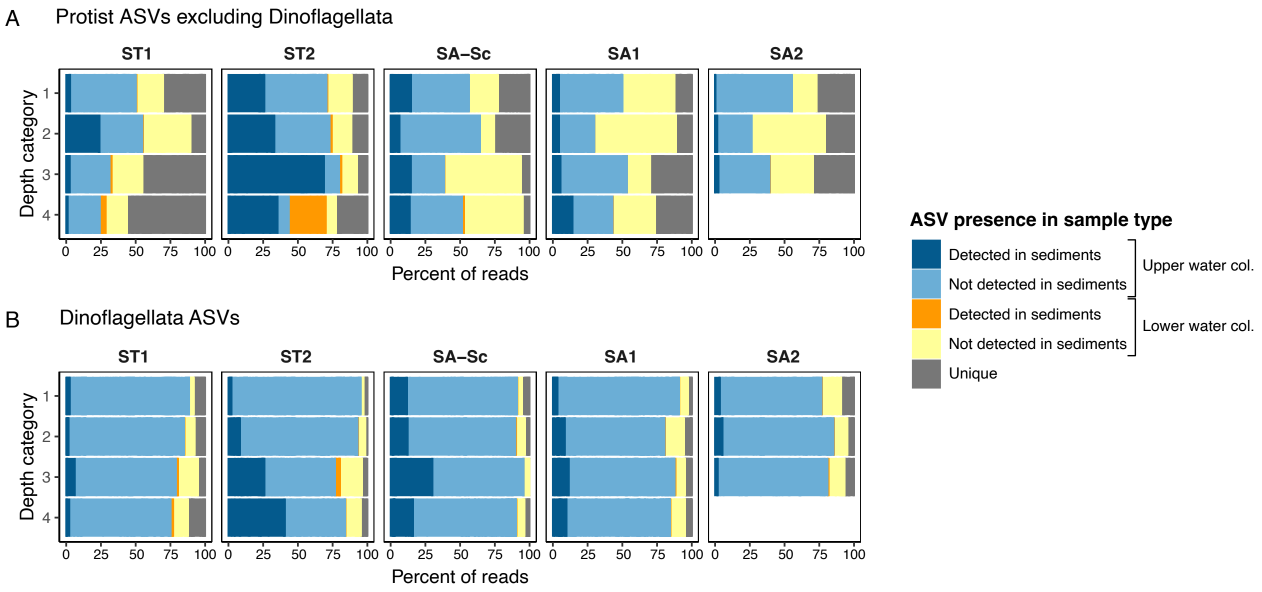


**Figure S6:** The relative abundance of (A) protist ASVs excluding Dinoflagellata and (B) Dinoflagellata ASVs in fixed trap samples that were shared between the upper or lower water column samples, further divided into ASVs that were detected or not detected in the sediments. Samples are grouped based on cycle and depth level sampled, corresponding to the depths specified in Table 1.

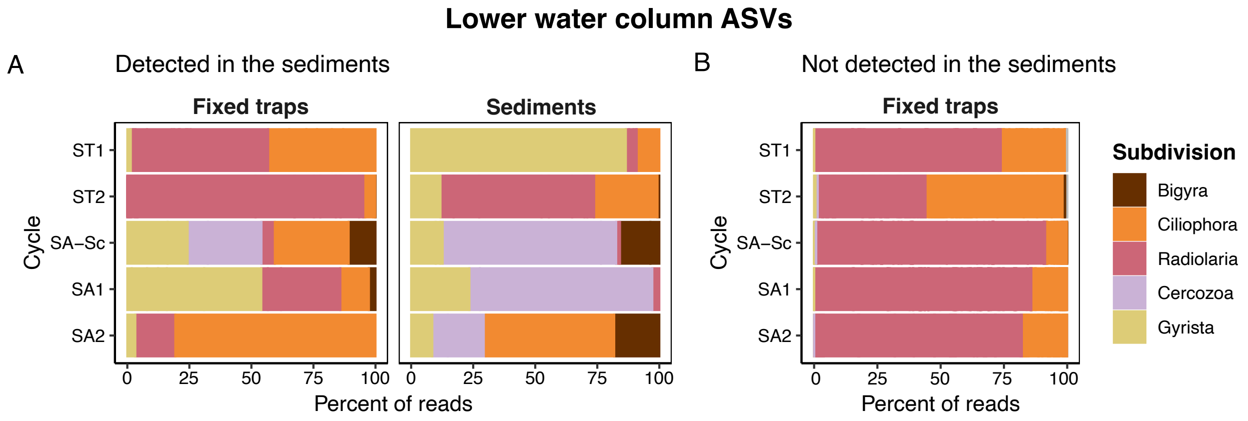


**Figure S7:** The relative abundance of protist ASVs from the lower water column samples at subdivision level (excluding Dinoflagellata) in fixed traps and sediment samples that were (A) detected in the fixed traps and sediment samples (represented by orange in Figure 6A), or (B) only detected in the fixed traps samples (yellow in Figure 6A).

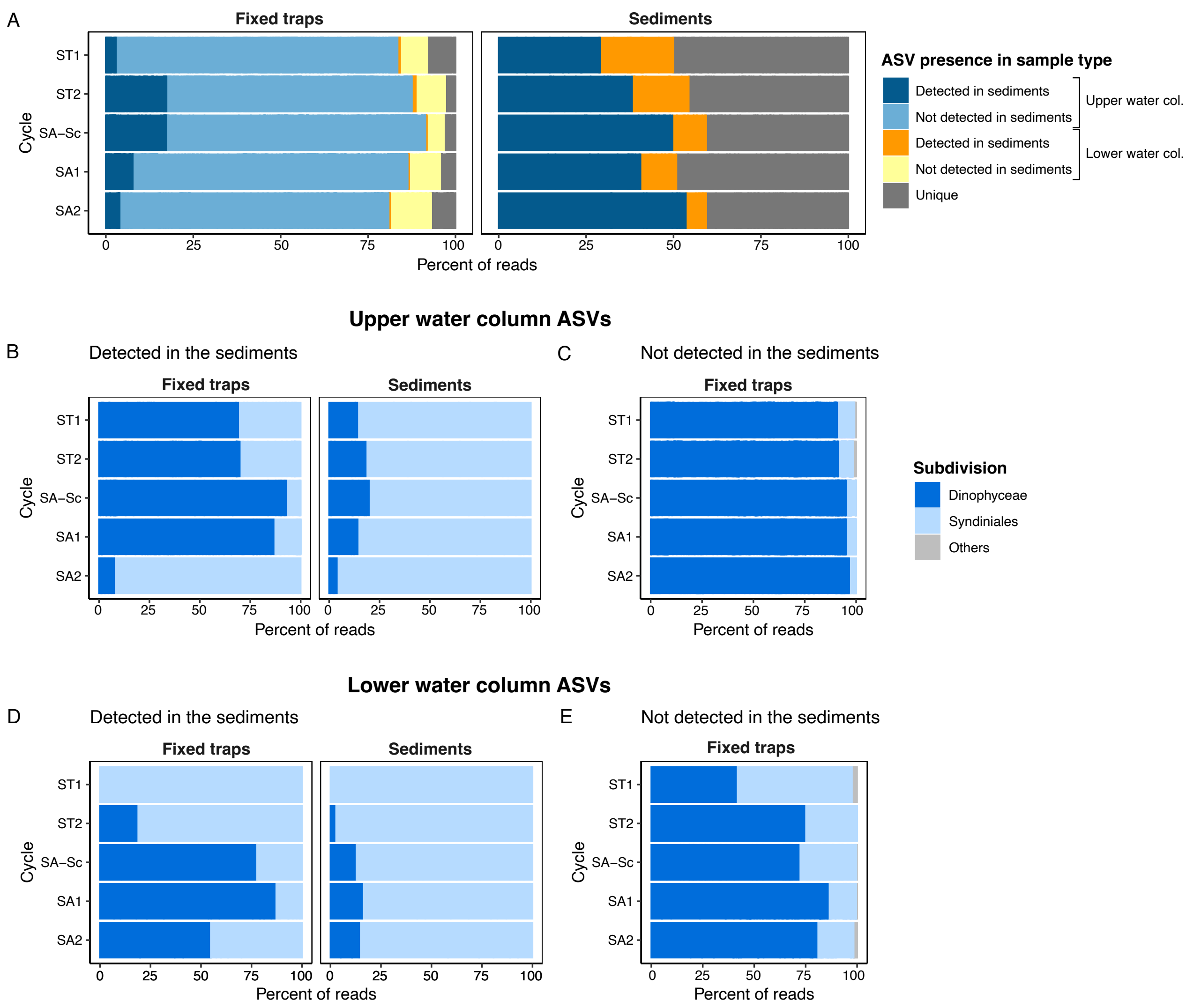


**Figure S8:** Subdivision Dinoflagellata from the fixed traps and sediment samples that were common between the upper and lower water column. A) Dinoflagellata ASVs in fixed traps and sediment samples were characterised as originating from the upper water column, further divided into ASVs exported or not exported to the sediments, or from the lower water column. Amongst ASVs from the upper water column, the relative abundance of Dinoflagellata at class level are presented for ASVs that were B) detected and C) not detected in the sediments (represented by dark blue and light blue in A, respectively). D) The relative abundance of Dinoflagellata ASVs at class level, from the lower water column in fixed trap and sediment samples (represented by yellow in A). ASVs from subtropical and subantarctic water masses were analysed separately.

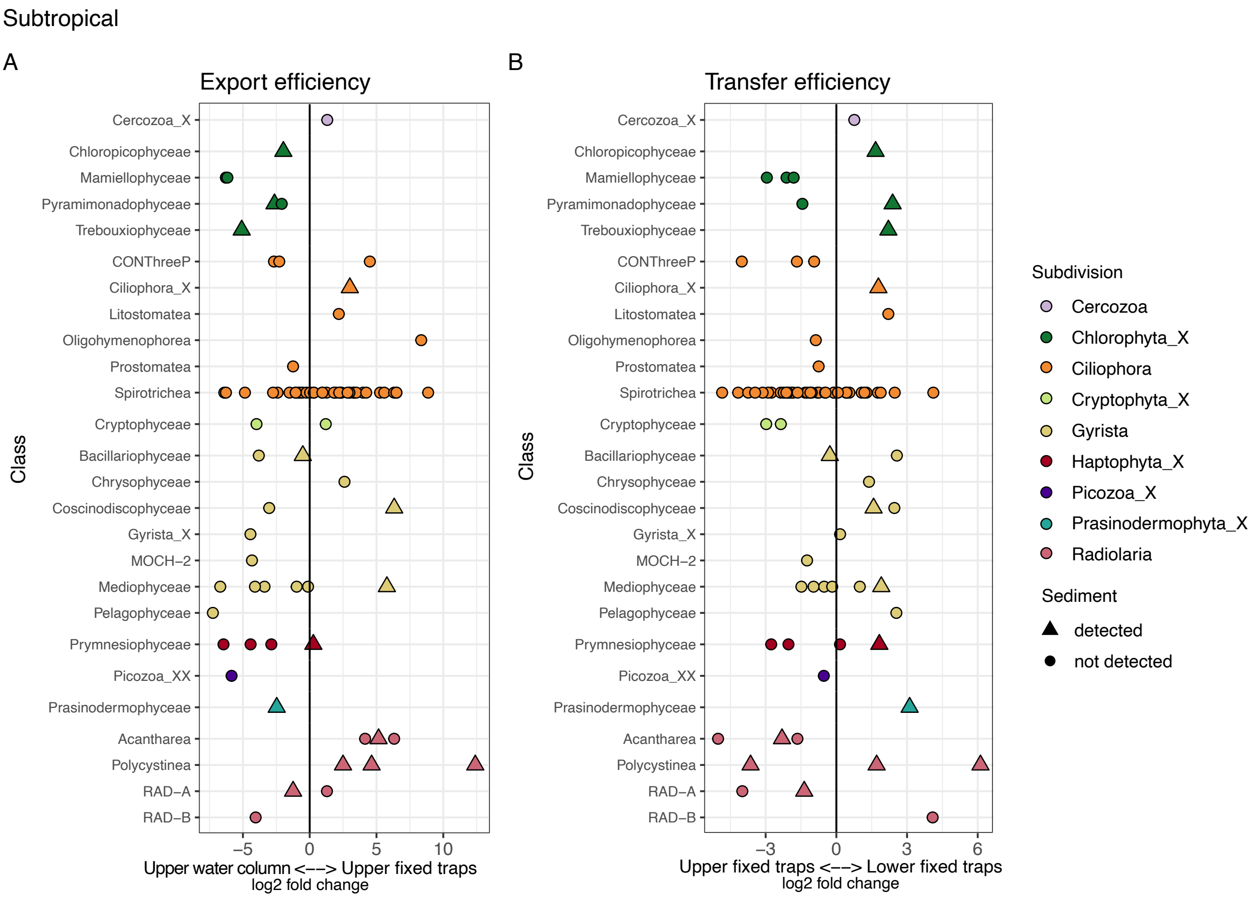


**Figure S9:** Differential gene abundance analysis (DESeq2) of protist ASVs (**excluding Dinoflagellata**) in subtropical cycles for A) between the upper water column and upper fixed traps (depth level 1 and 2) to show export potential, and B) between the upper fixed traps and lower fixed traps to show transfer potential. Colours represent subdivision of ASVs and shapes represent whether the ASV was detected in sediment of subtropical cycles.

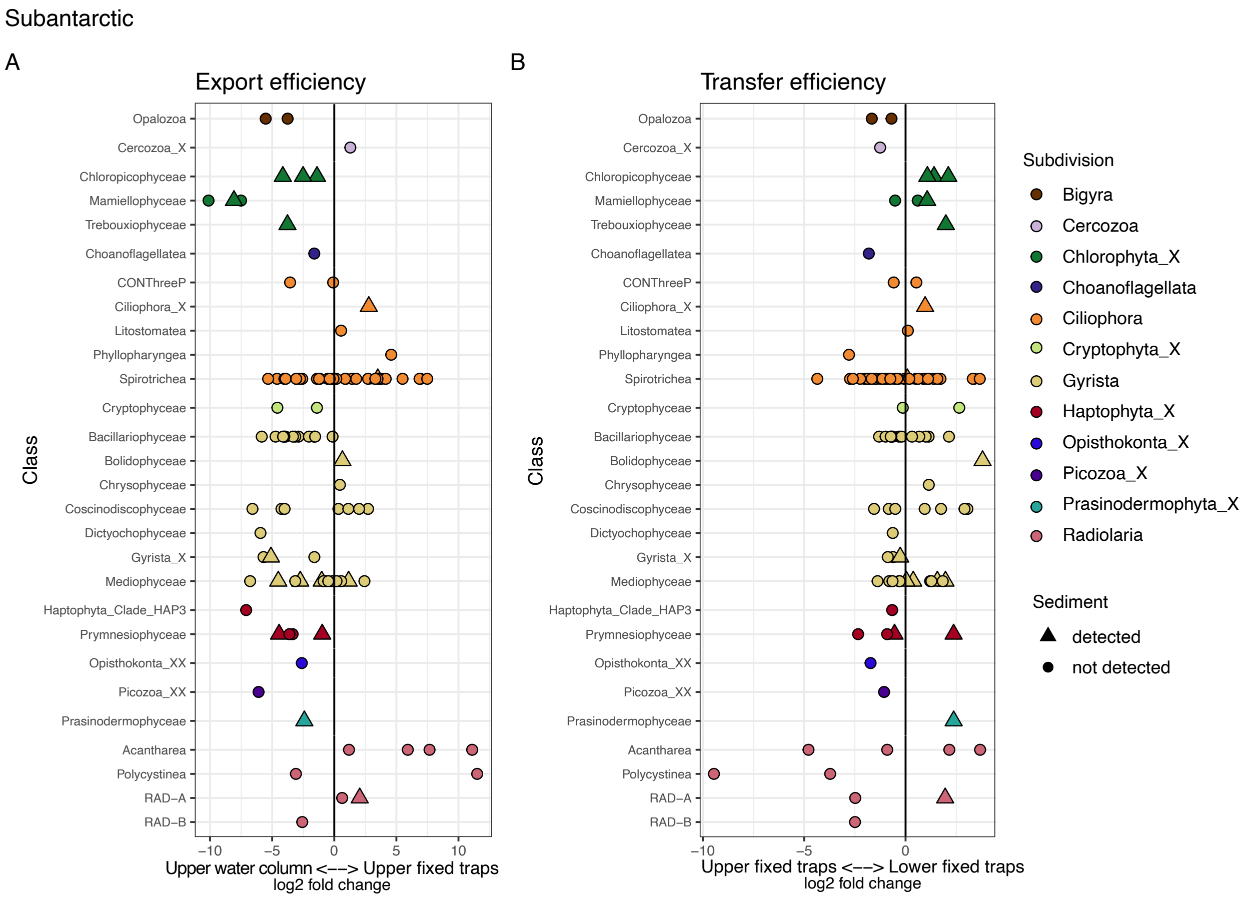


**Figure S10:** Differential gene abundance analysis (DESeq2) of protist ASVs (**excluding Dinoflagellata**) in subantarctic cycles for A) between the upper water column and upper fixed traps (depth level 1 and 2) to show export potential, and B) between the upper fixed traps and lower fixed traps to show transfer potential. Colours represent subdivision of ASVs and shapes represent whether the ASV was detected in sediment of subantarctic cycles.

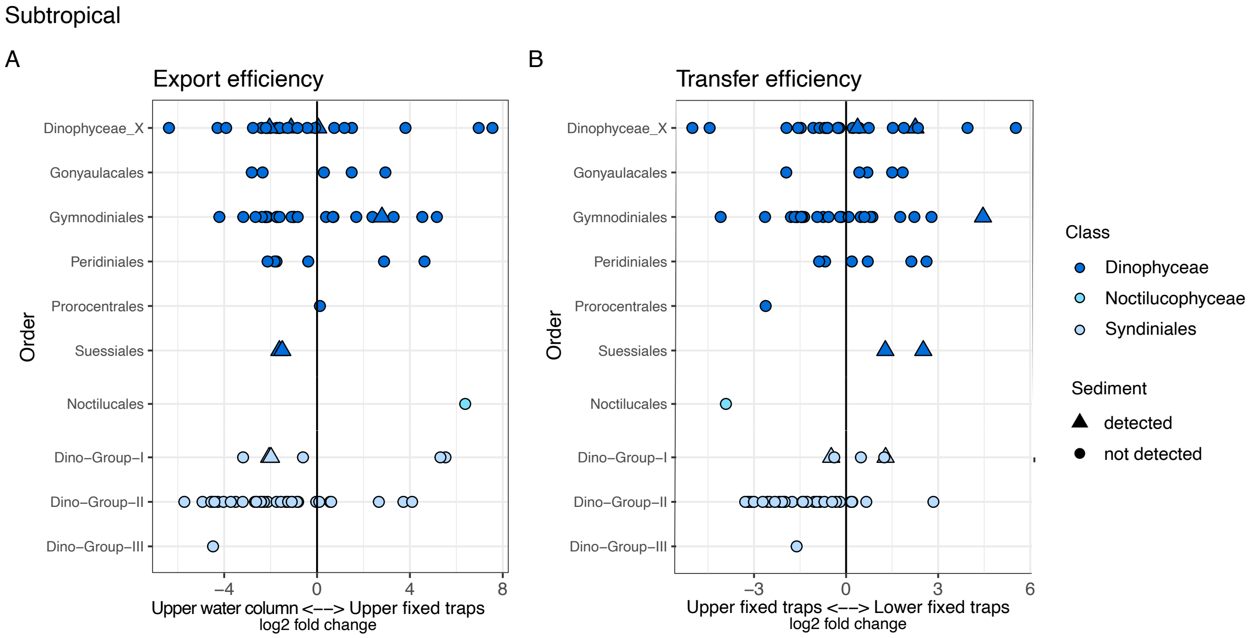


**Figure S11:** Differential gene abundance analysis (DESeq2) of **Dinoflagellata** ASVs in subtropical cycles for A) between the upper water column and upper fixed traps (depth level 1 and 2) to show export potential, and B) between the upper fixed traps and lower fixed traps to show transfer potential. Colours represent subdivision of ASVs and shapes represent whether the ASV was detected in sediment of subtropical cycles.

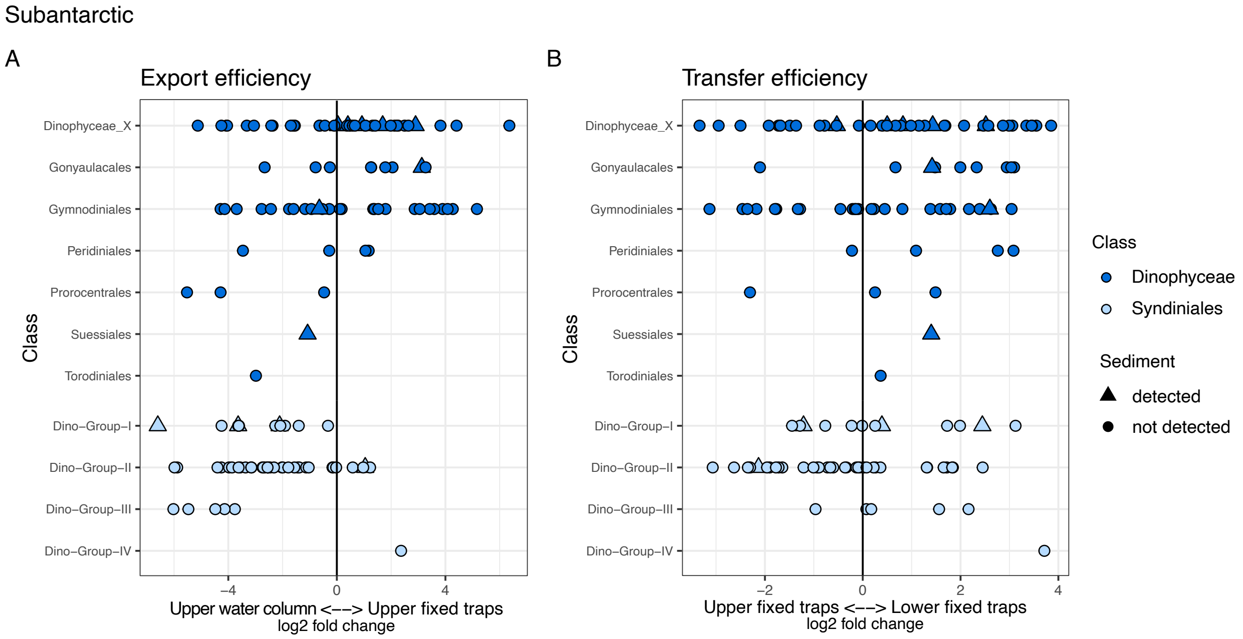


**Figure S12:** Differential gene abundance analysis (DESeq2) of **Dinoflagellata** ASVs in subantarctic cycles for A) between the upper water column and upper fixed traps (depth level 1 and 2) to show export potential, and B) between the upper fixed traps and lower fixed traps to show transfer potential. Colours represent subdivision of ASVs and shapes represent whether the ASV was detected in sediment of subantarctic cycles.
